# Integrative analysis identifies LATS1 as a context-specific regulator of melanoma progression and immune response

**DOI:** 10.64898/2026.11.12.766289

**Authors:** Nourhan Aboud, Donagh Egan, Andrew P Badrock, Pamela Kelly, Susan Peters, Brian Cloak, Kieran Wynne, Elena Merino Tejero, Sabina Susanu, Elaine O’Boyle, Luis Iglesias Martínez, Walter Kolch, Melinda Halasz, Adam Hurlstone, Lucía García-Gutiérrez, David Matallanas

## Abstract

Malignant melanoma incidence is increasing, and survival remains poor following metastasis. The main driver oncogenes are BRAF and NRAS. Advancements in targeted agents and immunotherapies have improved survival rates, especially for patients with BRAF mutant melanoma. However, no targeted therapy options are available for patients with NRAS-mutant melanoma. Unfortunately, the development of resistance to all existing therapies remains a challenge. Recent evidence has shown that the MST2/Hippo pathway is deregulated in melanoma. The MST2 pathway core kinase LATS1 is a signalling hub that mediates the crosstalk between cell death and proliferation networks, and its inactivation has been related to melanoma progression and resistance to targeted therapy. However, the mechanisms by which LATS1 influences melanoma biology remain unclear. Here, to investigate this role of LATS1, we developed a *lats1* knockout zebrafish melanoma model driven by either BRAF or NRAS mutations. Loss of Lats1 associates with reduced tumour-free survival in BRAF-mutant melanoma and a more invasive phenotype in NRAS-mutant melanoma, confirming a tumour-suppressive function. Integrating transcriptomics and proteomics datasets, we mapped molecular mechanisms mediated by Lats1 in tumours driven by these oncogenes, including pathways associated with the immune response regulated by Lats1. Subsequent validation in patient data confirmed an immune-evasive phenotype, associated with decreased LATS1 expression in BRAF-mutant melanoma. Additionally, the loss of LATS1 in NRAS mutant melanoma patients altered the expression of key DNA damage repair genes. Thus, this study identifies novel pathways controlled by LATS1 in melanoma, which aid in our understanding of the disease and offers new insights to be explored for the development of novel treatment strategies.

## Introduction

Cutaneous melanoma is the deadliest form of skin cancer, with rising incidence rates (1). Mutations in BRAF and NRAS drive approximately 50% and 20% of cases, respectively (2). While RAF inhibitors and immunotherapies have improved survival, most patients develop resistance within a year, and immunotherapy benefits only ∼50% of patients, with no predictive biomarkers and frequent adverse effects (2). Thus, deeper insights into melanoma biology and resistance mechanisms are essential.

Resistance to therapy often arises from signalling rewiring driven by oncogenic mutations (3). RAF kinases, including mutant BRAF, inhibit apoptotic signalling through kinase-independent mechanisms, notably by suppressing the MST2 pathway (4,5). This pathway, (6) includes mammalian MST1/2 and LATS1/2, which control cell fate and are frequently dysregulated in cancer (7). While much attention has focused on the MST1/2 effector YAP, the upstream kinase LATS1 remains largely underexplored in melanoma. Recent work from our group and others using melanoma cell models, murine models and clinical data supports a tumour-suppressive and pro-apoptotic role for LATS1 and the MST2 pathway in melanoma, both YAP-dependent and independent (8–12). Additionally, other evidence also implicate LATS1 in modulating the tumour immune microenvironment (13,14) suggesting broader functions in tumour biology.

In light of all this evidence, we hypothesised that the loss of LATS1 function disrupts key pro-apoptotic signalling, thereby contributing to melanoma development and progression. To test this hypothesis and delineate LATS1-dependent signalling modules in melanoma, we aimed to characterise the phenotypes associated with LATS1 loss *in vivo*. We developed mutant NRAS and BRAF-driven zebrafish melanoma models on the genetic background of a *lats1* knockout. Zebrafish offer substantial advantages for studying this cancer type due to significant genetic homology with humans, with over 80% of human disease-related genes conserved (15), the capacity to assess large numbers of animals for tumour burden in a cost-effective manner, and an intact immune system not present in widely used mouse xenograft models (15,16). Moreover, it has been shown that zebrafish melanoma models driven by melanocyte-specific expression of mutant BRAF or NRAS in a *tp53*-mutant background recapitulate human melanoma pathology (15).

Here, we report that *lats1* plays distinct roles in melanoma depending on the oncogenic driver. In BRAF^V600E^ melanoma, *lats1* loss reduced tumour-free survival and promoted an immune-evasive phenotype. In NRAS^Q61L^ melanoma, *lats1* loss was associated with increased invasive expression signatures and altered DNA repair signalling genes. These findings were validated in human datasets, where *LATS1* deletions in BRAF-mutant melanoma correlated with reduced IFN-γ signalling and immune cell infiltration. Altogether, our results identify LATS1 as a context-specific tumour suppressor with potential implications for immunotherapy response and therapeutic targeting in melanoma.

## Results

### *lats1* expression results in differential tumour-free survival in BRAF^V600E^ but not NRAS^Q61L^ melanoma

To investigate LATS1’s role in melanoma development *in vivo*, we used the *tp53^M214K^; mitfa^w2^* zebrafish model and generated a *lats1* knockout line. In this model, Mitfa loss blocks melanocyte development, as Mitfa is the key transcriptional regulator of this process (17). The miniCoopR system restores Mitfa expression and melanocyte development, while enabling expression of genes of interest under the *mitfa* promoter, restricting their expression to melanocytes. Impaired p53 function allows melanoma induction when either BRAF^V600E^ or NRAS^Q61L^ are expressed via the miniCoopR system (17). We created a *lats1* knockout line on the *tp53^M214K^; mitfa^w2^* background using CRISPR/Cas9, generating a 52 bp deletion that caused a frameshift within the *lats1* gene and consequently its loss of expression (Fig. 1A, Supplementary Fig. S1A). To induce melanoma, one-cell stage embryos were injected with miniCoopR constructs carrying human BRAF^V600E^ or NRAS^Q61L^, and larvae with comparable melanocyte rescue were raised to adulthood (Fig. 1B). A GFP-expressing miniCoopR plasmid served as control. Protein expression of MST2 pathway components and oncogenes was validated by western blot (Supplementary Fig. S1B, C). Transgenic lines and animal numbers are detailed in Supplementary Fig. S1D.

**Figure 1.**
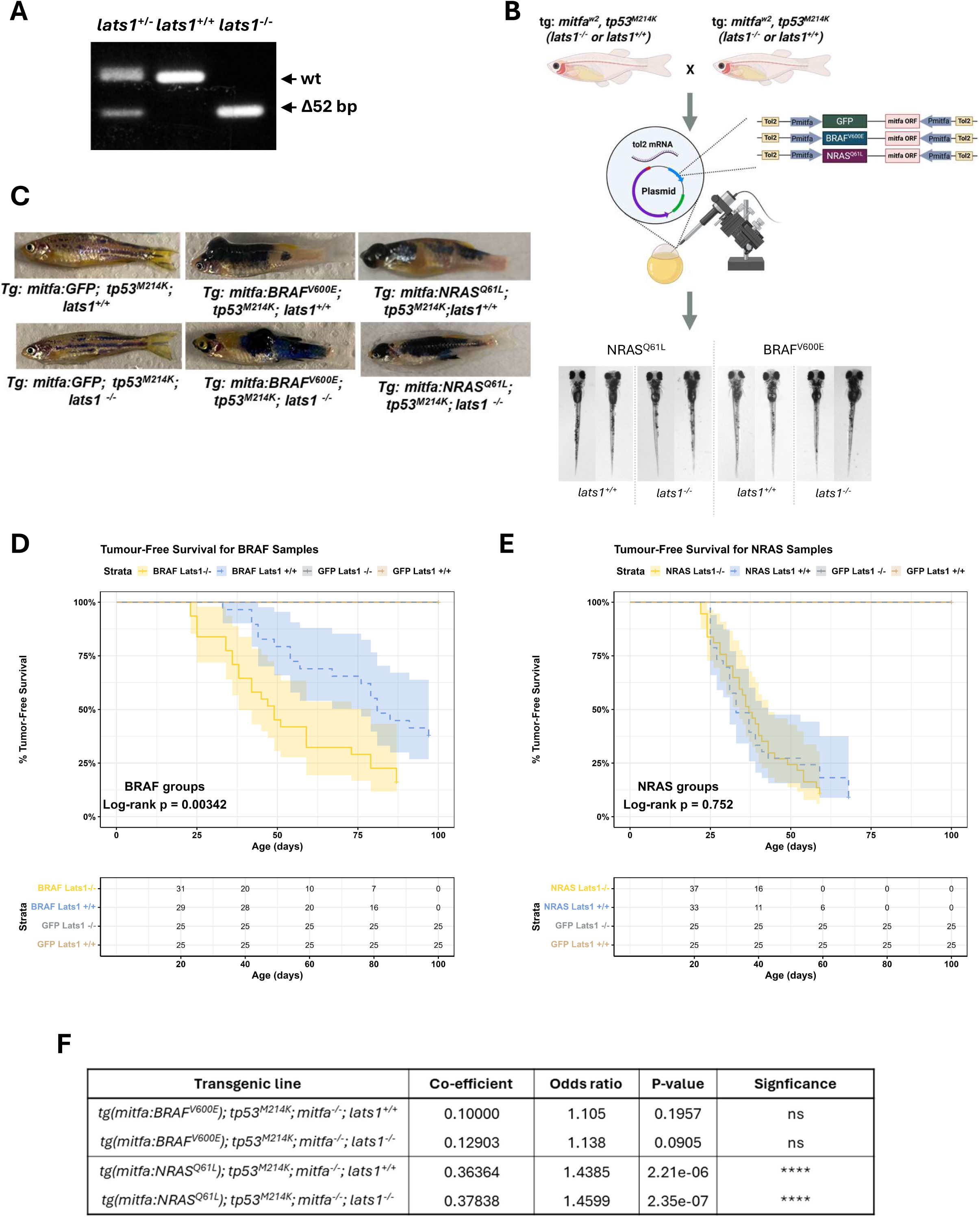
Generation of a Lats1 -/- zebrafish melanoma model driven by BRAF and NRAS mutations. A) PCR agarose gel confirming the *lats1* knockout in zebrafish embryos. B) Schematic showing the microinjection of zebrafish embryos for the induction of melanocyte development with the desired genetic background using MiniCoopR plasmids and images of the resulting embryos with melanocyte rescue. C) Representative images of adult zebrafish post-microinjection showing the melanocyte rescue in the control groups and the melanoma development in the experimental groups. (D-E) Kaplan Meier curves showing the percentage of tumour-free survival for zebrafish harbouring BRAF^V600E^ melanoma (D) and NRAS^Q61L^ melanoma (E) the associated risk tables. Statistical analysis between the experimental groups with regards to the status of *lats1* expression was performed using the Log-rank test. F) Table showing results of the logistic regression analysis with the odds ratio of developing multiple tumours for each genetic background and the corresponding p-value.

As a first step to determine the possible role of *lats1* in melanoma development, we focused on the study of the macroscopic phenotypes in these animals. Both *tp53^M214K^; lats1^+/+^ and tp53^M214K^; lats1^-/-^* GFP-control groups showed normal pigmentation and no nevi or tumours, indicating that neither *mitfa* expression nor *lats1* loss alone triggers spontaneous tumorigenesis in this model (Fig. 1C). In contrast, animals injected with BRAF^V600E^ or NRAS^Q61L^ miniCoopR plasmids developed diffuse black patches reassembled nevi (Fig. 1C). NRAS^Q61L^-expressing animals developed protruding tumours in 3-4 weeks, while BRAF^V600E^ fish showed tumour onset at 7–8 weeks (Fig. 1C, D C E). Remarkably, *lats1* deletion accelerated tumour onset in BRAF^V600E^ but not NRAS^Q61L^ fish. Kaplan-Meier analysis confirmed that *lats1* loss significantly reduced tumour-free survival in BRAF^V600E^ fish (Log-rank p=0.0034; Fig. 1D), suggesting a protective role for *lats1* in BRAF-driven melanoma.

Interestingly, we found that melanoma driven by an NRAS mutation correlated with reduced tumour-free survival compared to BRAF mutant melanoma, indicating that the zebrafish NRAS tumours might be more aggressive, in line with some observations in human melanoma (18) (Fig. 1D C E). In this NRAS^Q61L^ fish line, tumours appeared at 25 days post-injection when *lats1* is expressed, and at 22 days in *lats1^-/-^* animals (Fig. 1E). However, *lats1* loss had only a modest effect on tumour onset and did not significantly alter tumour-free survival (Log-rank p=0.75; Fig. 1E), indicating that *lats1* does not influence NRAS-driven melanoma progression.

As an additional parameter to characterise phenotypic differences driven by the two oncogenes and the impact of *lats1* loss, we quantified the number of tumours per fish in each group. NRAS^Q61L^ fish developed more tumours than BRAF^V600E^ fish, confirming greater aggressiveness (Supplementary Fig. S1E). Regression analysis linked multiple tumour development to NRAS^Q61L^ expression (Fig. 1F). *lats1* loss slightly increased the odds of multiple tumours (from 43.81% to 45.99%), but this was not statistically significant (z = -0.146, p = 0.884). Overall, our data show that BRAF^V600E^ and NRAS^Q61L^ mutations drive melanoma differently in zebrafish, and that *lats1* plays a tumour-suppressive role specifically in BRAF-driven melanoma.

### Histological analysis reveals a role for *lats1* in promoting more invasive phenotypes in BRAF^V600E^ and NRAS^Q61L^ melanoma

The histological resemblance between human and zebrafish melanoma is well established (15). Therefore, as a second step of our analysis, we investigated the effects of *lats1* expression in melanoma at the microscopic level to elucidate the potential role of this tumour suppressor in tumour morphology. To do this, we analysed the impact of *lats1* expression on the histological organisation of zebrafish melanoma using parameters analogous to those assessed in human melanoma patients. Specifically, we determined the depth of melanoma invasion and also measured zebrafish skin thickness, which would be akin to the Breslow depth, a standard criterion used in melanoma staging (19). Analysis of transverse zebrafish sections from the BRAF^V600E^ groups revealed distinct invasion patterns based on the *lats1* status. The BRAF^V600E^*; lats1^+/+^* group, compared to the *lats1^-/-^* group, exhibited more superficial tumour development, with pigmentation predominantly localised to the skin and adjacent muscle (Fig. 2A, B and Supplementary Table 1). Importantly, evidence of melanoma muscle invasion was observed in only one of three BRAF^V600E^*; lats1^+/+^* group compared to all four fish in the BRAF^V600E^*; lats1^-/-^* samples (Supplementary Table 1). Furthermore, none of the BRAF^V600E^*; lats1^+/+^* fish showed pigmented or neoplastic cells invading the spinal cord in striking contrast to the BRAF^V600E^*; lats1^-/-^* group where 3 out of 4 fish demonstrated a more invasive phenotype with pigmented cells invading through the muscle to the spinal cord (Fig. 2B and Supplementary Table 1). In contrast, all NRAS^Q61L^ fish, regardless of *lats1* status, showed melanoma invasion through the muscle layer (Fig. 2C, D and Supplementary Table 1). Notably, we detected the presence of melanoma cells in the brain in 3 out of 5 fish in the NRAS^Q61L^*; lats1^-/-^* group, whereas none were detected in the NRAS^Q61L^*; lats1^+/+^* or any of the BRAF^V600E^ groups (Fig. 2E and Supplementary Table 1). Our model does not allow us to determine if these brain tumoral cells originate in brain cells or are the result of metastatic cells forming secondary tumours. Nevertheless, given that brain metastasis develops in 50% of melanoma patients with advanced-stage disease and correlates with poorer prognosis (20), this finding highlights the need to further explore LATS1’s role in NRAS-driven melanoma progression. Additionally, during histological analysis, we noted differences in skin thickness across genetic groups. Since depth of melanoma invasion and Breslow depths are standard criteria in melanoma staging (19), we measured the thickest portion of the skin/epithelial layer in the dorsal region above the dorsal spinous process and just rostral to the dorsal fin in transverse sections (Supplementary Fig. S2A). Amongst the four melanoma groups, the NRAS^Q61L^*; lats1^-/-^* samples developed a significantly thicker skin layer compared to the NRAS^Q61L^*; lats1^+/+^* samples (p= 0.037), as well as the BRAF^V600E^*; lats1^+/+^* (p=0.0048) and the BRAF^V600E^*; lats1^-/-^* (p=0.015) (Fig. 2F). In contrast, we did not find a significant change in skin thickness when comparing the BRAF^V600E^ samples with regards to the status of *lats1* expression (p=0.45). Collectively, these histological findings support a role for *lats1* in modulating melanoma invasiveness, with effects varying according to the oncogenic driver.

**Figure 2.**
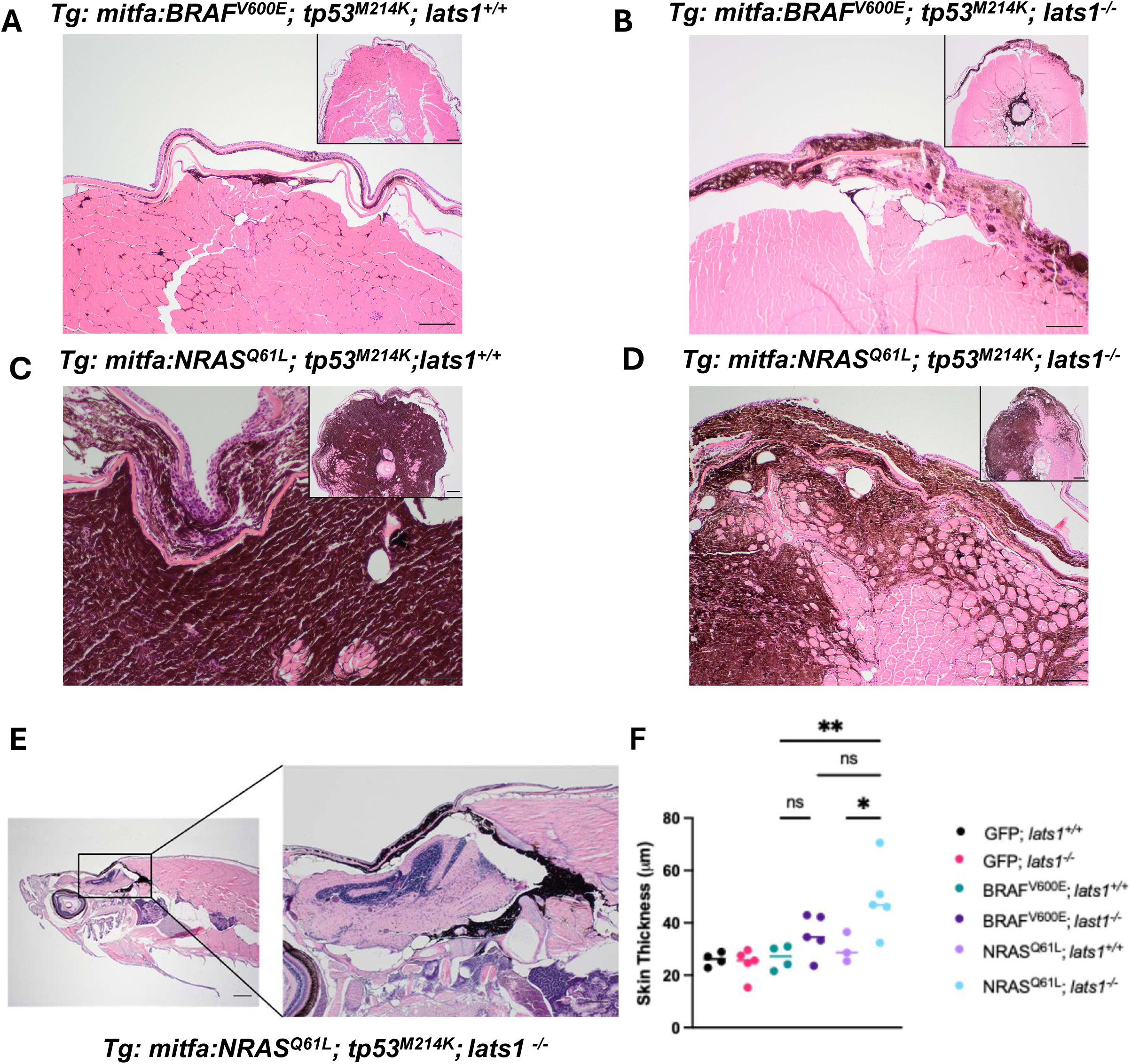
Histological analysis of zebrafish sections reveals a more aggressive phenotype with lats1 loss. (A-D) Representative transverse sections stained with haematoxylin and eosin of zebrafish showing muscle and spinal cord melanoma infiltration (4X) and the magnification of the skin layer in each section (10X) for BRAF^V600E^; *lats1^+/+^* (A), BRAF^V600E^; *lats1^-/-^* (B), NRAS^Q61L^; *lats1^+/+^* (C) and NRAS^Q61L^; *lats1^-/-^* (D) groups. The arrows indicate evidence of tumour invasion. E) Representative longitudinal section stained with haematoxylin and eosin for NRAS^Q61L^; *lats1^-/-^*^-^ (4X) showing melanoma brain invasion (10X). F) Scatter plot showing the thickness of the skin layer measured in µm of zebrafish in the different genetic groups. Statistical analysis: Ordinary one-way ANOVA. *p<0.05, **p<0.01.

### *lats1* regulates immune-related pathways, and its loss promotes pro-survival signalling in BRAF^V600E^ melanoma

The previous data indicated that *lats1* plays a key role in melanoma development in zebrafish. Based on our hypothesis and prior findings (12), the more aggressive phenotype in *lats1* knockout tumours likely results from deregulated signalling networks normally controlled by this tumour suppressor. To explore this, we characterised the molecular changes associated with *lats1* deletion using a systematic, unbiased approach. To get in-depth mechanistic insight, we generated high-throughput transcriptomics and proteomics datasets from zebrafish superficial tumours and reconstructed signalling pathways computationally using the individual datasets, followed by integrative analysis. Our approach avoids the limitations that the individual use of this type of dataset has, including low mRNA-protein correlation (21). Tumours were surgically excised and split for RNA and protein extraction, producing matched transcriptomic and proteomic datasets (Fig. 3A, B). Quality control confirmed high coverage of the zebrafish transcriptome and proteome (Fig. S3A, B). Differential expression analysis followed by gene set pathway enrichment (GSEA) using Gene Ontology (GO) of the RNA-seq data showed that the top five pathways positively enriched in BRAF^V600E^*; lats1^+/+^* versus *lats1^-/-^* were immune-related, suggesting a role for Lats1 in regulating the immune landscape in melanoma (Fig. 3C). In total, 14 immune-related pathways were significantly enriched in the *lats1^+/+^* group, with overlapping genes (Supplementary Fig. 3C). A heatmap of genes shared across five or more GO terms clustered most BRAF^V600E^ samples by *lats1* status (Fig. 3D). The transcriptional changes comprise both tumour-intrinsic inflammatory signalling and signatures compatible with immune-cell infiltration/activation. Upregulated genes in the *lats1^+/+^* group included interferon signalling components (*stat1b, ifng1*, *irf1b*, *ifit* family) and the MHC-I gene *b2m*, consistent with activation of the IFN response program in melanoma cells. In parallel, the increased expression of immune cell-associated genes including the B-cell markers (*cd180*, *cd40*), the cytotoxic effector gene perforin-related *prf1.1*, and the inflammatory nitric oxide synthase genes *nos2a* and *nosa2b* (Fig. 3D) indicate enhanced immune cell infiltration and activation within the tumour microenvironment. Altogether, these findings suggest the involvement of *lats1* in promoting an immune response in our BRAF^V600E^ melanoma model.

**Figure 3.**
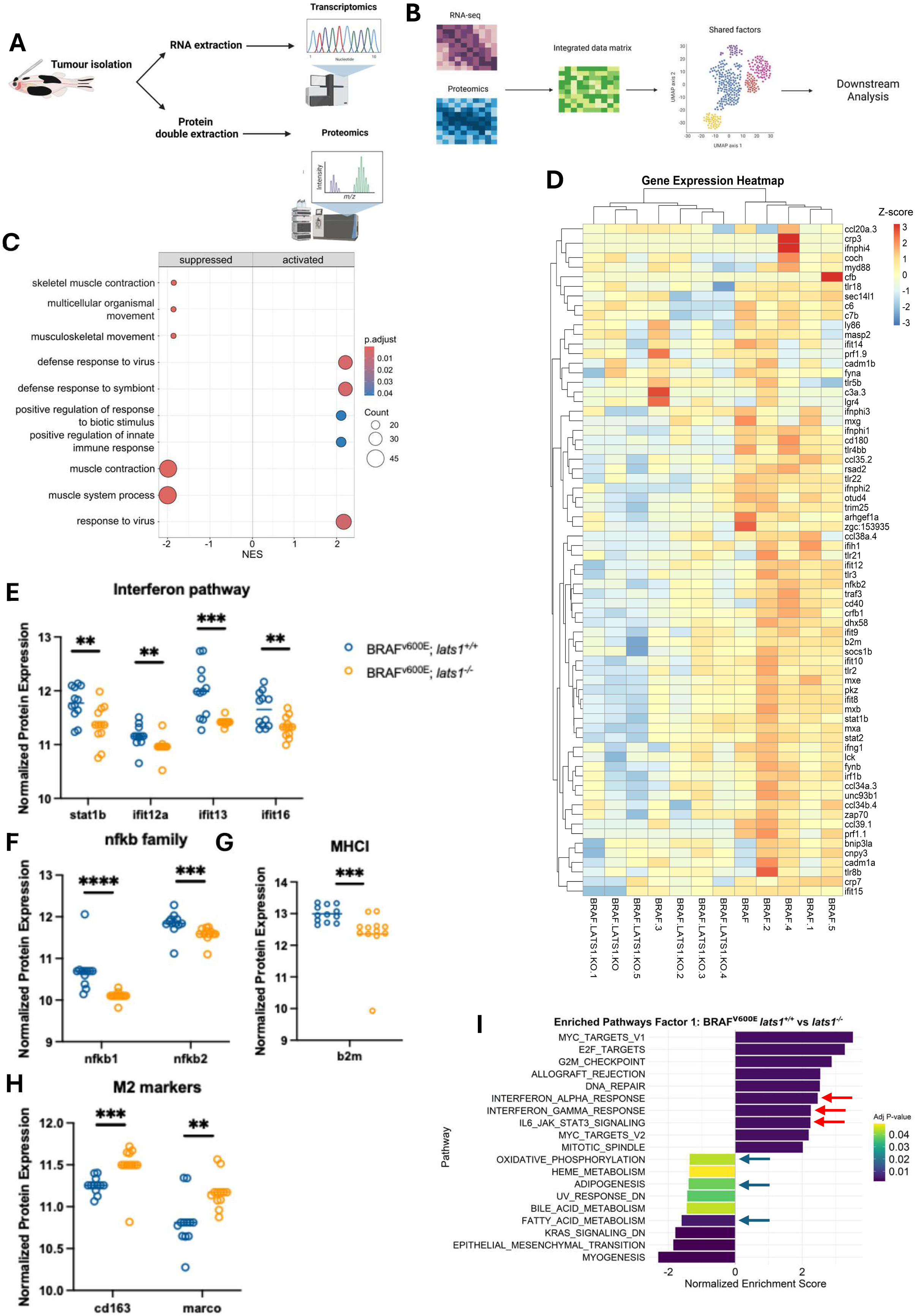
Pathway enrichment analysis for BRAF^V600E^ zebrafish melanoma correlate lats1 expression with immune activation. A) Simplified schematic showing the work flow to generate matching transcriptomics and proteomics dataset from zebrafish tumours. B) Workflow for integrating the multi-omics datasets to generate shared sources of variations called factors. C) Bubble plot illustrating the results of Gene set enrichment analysis (GSEA) for Gene Ontology Biological Processes (GO BP) derived from RNA-seq data. The plot shows the top 5 significantly activated and suppressed pathways in BRAF^V600E^; *lats1^+/+^* compared to *lats1^-/-^*.The colour of the bubble corresponds to the p-adjusted values and the size of the bubble corresponds to the number of genes contributing to each pathway. D) Heatmap showing the expression of overlapping genes obtained from the pathway similarity network across the BRAF^V600E^; *lats1^+/+^* (BRAF – BRAF.5) and *lats1^-/-^*. (BRAF.LATS1.KO – BRAF.LATS1.KO.5) tumours. (E-H) Scatter plots showing the normalized expression for protein markers of the interferon pathway (E), NF-κB pathway (F), Major histocompatibility I (G) and Macrophage M2 phenotype (H) in BRAF^V600E^; *lats1^+/+^* and *lats1^-/-^* samples. Shapiro-Wilk test was used to determine normal distribution. For normally distributed data statistical significance was calculated using an Unpaired t-test and for non-normally distributed data significance was calculated using the Mann-Whitney test. **p<0.01, ***p<0.001, ****p<0.0001. I) Bi-directional bar plots showing the normalized enrichment score (NES) and the adjusted p-value obtained from the GSEA Hallmarks database analysis for the genes identified in factor 1.

This was further supported by mining the proteomics dataset which replicated the increased levels of the interferon pathway members Stat1b, Ifit12a, Ifit13a and Ifit16 on the protein levels in the BRAF^V600E^*; lats1^+/+^* versus *lats1^-/-^* (Fig. 3E). Higher expression of the nuclear factor-kB (NF-kB) pathway proteins Nfkb1 and Nfkb2 were also observed in the *lats1^+/+^* compared to *lats1^-/-^* samples (Fig. 3F). These transcription factors are established mediators of the inflammatory response and immune cells activation (22). Additionally, further supporting our RNA-seq findings, the MHC-I protein levels of B2m were significantly increased in the *lats1^+/+^* samples (Fig. 3G). This was concurrent with decreased levels of the tumour-promoting M2 macrophage markers Cd163 and Marco (Fig. 3H).

Finally, we used the Multiset correlation and factor analysis (MCFA) method (23) to integrate both transcriptomics and proteomics datasets and identify shared sources of variation across both datasets, termed factors. The primary source of variation between the BRAF^V600E^*; lats1^+/+^* versus *lats1^-/-^* tumours were defined by the set of genes in factor 1 (p=0.019) (Supplementary Fig. S3D). GSEA using the Hallmarks of Cancer database revealed enrichment of interferon alpha and gamma responses and IL-6 JAK/STAT signalling in the *lats1*^+/+^ samples (Fig. 3I), consistent with an immune responsive state (24). In contrast, oxidative phosphorylation, adipogenesis, and fatty acid metabolism were depleted in the *lats1^+/+^* tumours compared to the *lats1^-/-^*, indicating a shift towards lipid dependent oxidative metabolism upon *lats1* loss and an immune evasive state (25). These results support the hypothesis that *lats1* expression associates with an immunogenic phenotype, whereas its loss correlates with a shift towards oxidative metabolism and reduced immune activation.

### *lats1* regulates DNA repair and cell cycle progression genes in NRAS^Q61L^ melanoma

Next, to assess the molecular impact of *lats1* loss in NRAS^Q61L^-zebrafish melanoma, we followed the same workflow applied for our BRAF^V600E^ samples by exploring the individual RNA-seq and proteomics datasets, then performing data integration using MCFA. GSEA analysis using the KEGG database revealed several significantly enriched pathways, however in this case the negatively enriched, i.e. suppressed, pathways with *lats1* expression downstream the NRAS^Q61L^ mutation were particularly intriguing. Notably, we saw that genes related to DNA repair mechanisms are suppressed with *lats1* expression (Fig. 4A). More specifically, the Fanconi Anemia (FA) pathway was the highest negatively enriched in NRAS^Q61L^*; lats1^+/+^* compared to *lats1^-/-^* (Fig.4A). The FA pathway is well described to repair DNA crosslinks, maintain genome integrity and is reported to be upregulated in melanoma (26). Indeed, the expression levels of the core pathway components *rad51c, fancl, fancg and fance* were significantly downregulated in the NRAS^Q61L^*; lats1^+/+^* group (Fig. 4B). Consistently, we saw a negative enrichment of genes involved in both double strand breaks (DSBs) DNA repair mechanisms, Homologous recombination (HR) and Non-homologous end joining (NHEJ), and a concomitant suppression of genes involved in DNA replication (Fig. 4A).

**Figure 4.**
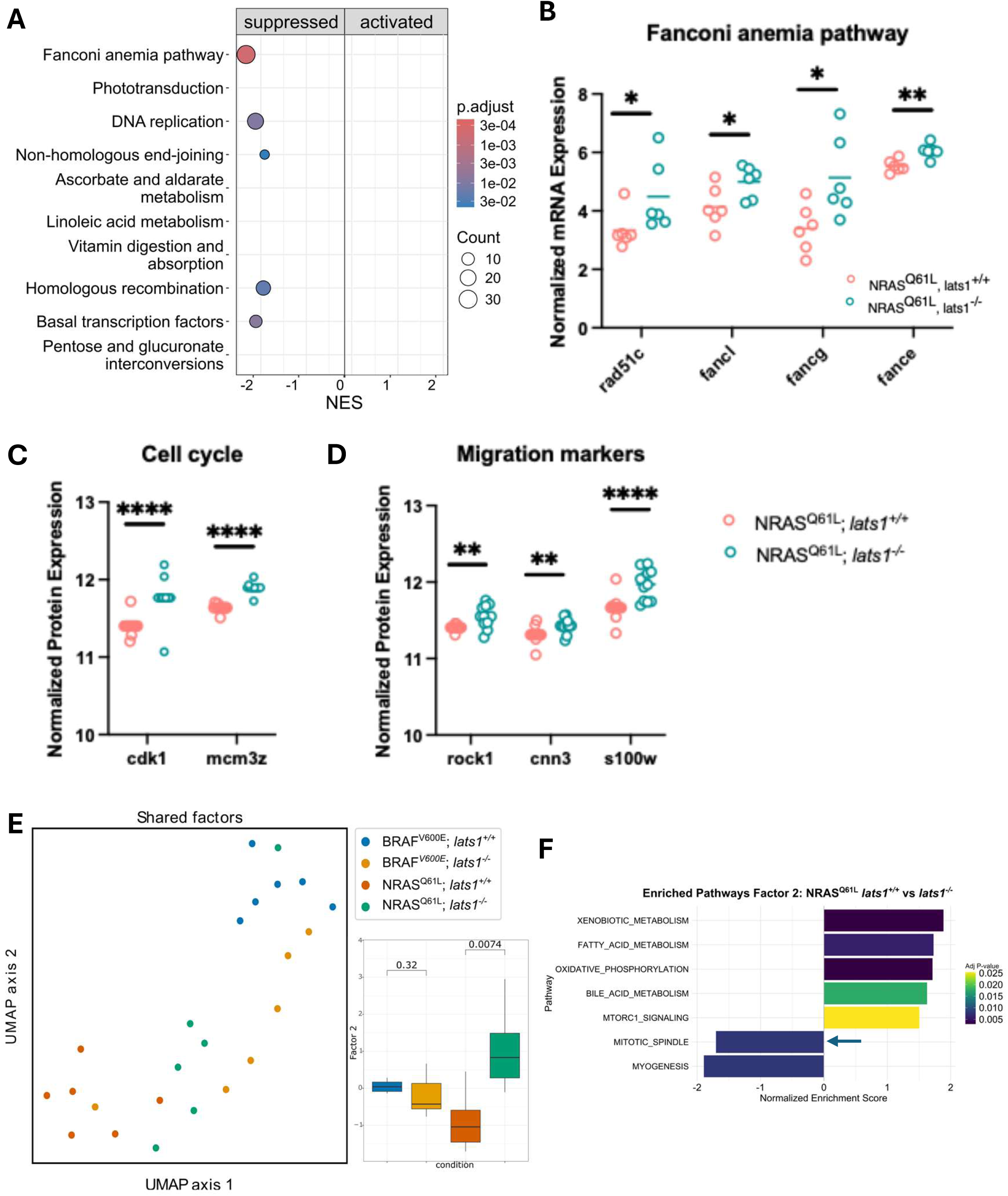
Pathway enrichment analysis for zebrafish NRAS^Q61L^ tumours associate lats1 expression with altered DNA repair mechanisms. A) Bubble plot illustrating the results of Gene set enrichment analysis (GSEA) using the KEGG database derived from RNA-seq data of NRAS mutant tumours. The plot shows the top 5 significantly activated and suppressed pathways in NRAS^Q61L^; *lats1^+/+^* compared to *lats1^-/-^*. The colour of the bubble corresponds to the p-adjusted values, and the size of the bubble corresponds to the number of genes contributing to each pathway. B) Box plots for expression values of significant genes contributing towards the enrichment of the Fanconi anaemia pathway in the NRAS^Q61L^; *lats1^+/+^* and *lats1^-/-^*groups. Normal distribution was assessed using the Shapiro-Wilk test, for normally distributed genes significance was calculated using an Unpaired t-test and for non-normally distributed genes significance was calculated using the Mann-Whitney test. *p< 0.05, **p<0.01. (C, D) Scatter plots of the normalized protein expression levels involved in Cell cycle (C) and Migration (D). Normal distribution was assessed using the Shapiro-Wilk test, for normally distributed genes significance was calculated using an Unpaired t-test and for non-normally distributed genes significance was calculated using the Mann-Whitney test. **p< 0.01, ***p<0.001, ****p<0.0001. E) UMAP showing the shared factors generated by multi-omics data integration and the clustering of the different samples with the associated box-plots showing the significance of factor 2 which is contributing towards the variances of the NRAS^Q61L^ melanoma groups. F) Bi-directional bar plots showing the normalized enrichment score (NES) and the adjusted p-value obtained from the GSEA Hallmarks database analysis for the genes identified in factor 2.

The previous data suggested that the loss of *lats1* expression in zebrafish NRAS^Q61L^ melanoma, conferred an advantage in melanoma DNA repair mechanisms and allowed tumour cells under stress and constant proliferation to overcome DNA damage leading to cell cycle progression. To explore our hypothesis further, we turned to our proteomics dataset for markers of cell cycle and tumour progression. Consistently, we found a significant downregulation in the protein levels of the cell cycle markers Cdk1 and Mcmz3 in NRAS^Q61L^*; lats1^+/+^* compared to *lats1^-/-^* (Fig. 4C). Additionally we found that with the loss of *lats1* there is a significant upregulation of Rock1, Cnn3 and S100w levels (Fig. 4D), all markers previously described to mediate melanoma invasion and migration (27). This is consistent with our earlier histological findings showing the increased invasiveness of melanoma cells in the NRAS^Q61L^*; lats1^-/-^* cohort. Therefore, our proteomics data further suggest that *lats1* loss of expression mediates a more aggressive melanoma phenotype downstream NRAS^Q61L^ in zebrafish.

Finally, we wanted to investigate if *lats1* has immune-related roles in NRAS^Q61L^ melanoma, similar to what we observed in the BRAF^V600E^ tumours. Therefore, we performed GSEA using GO and found only 3 immune-related pathways that were positively enriched in the NRAS^Q61L^*; lats1^+/+^* tumours compared to the *lats1^-/-^* group (Supplementary Fig. S4A). This shows that the effect of *lats1* on the anti-tumour immune response is limited in the case of NRAS^Q61L^ melanoma and its protective effect seems to be mostly modulated through the impediment of DNA repair mechanisms. Next, we applied the MCFA pipeline (23) to integrate the transcriptomics and proteomics datasets for the NRAS^Q61L^ melanoma zebrafish samples to have a wholesome overview of the molecular changes occurring between these two groups. The genes captured in Factor 2 separated our NRAS samples according to the status of *lats1* expression (Fig. 4E). GSEA of these genes using the Hallmarks of Cancer database supported our earlier findings showing a suppression of the mitotic spindle pathway with *lats1* expression (Fig. 4F). Further strengthening the link between *lats1* expression, genome stability and cell cycle progression. Finally, GSEA of the integrated data did not provide evidence regarding the involvement of *lats1* in an anti-tumour immune response in NRAS^Q61L^ tumours, confirming our previous results.

### Analysis of clinical datasets shows LATS1 has differential roles in melanoma which are dependent on the driver mutation

The zebrafish model resulted in insightful information about pathophysiological and molecular changes associated with *lats1* expression and the two main melanoma driver oncogenes. Therefore, it was important to further validate our findings and illustrate their relevance to the study of human melanoma by using publicly available patient data. To address this, we resorted to The Cancer Genome Atlas (TCGA) Firehose Legacy dataset (28). This included data from a total of 480 melanoma patients, of which there were 236 BRAF mutations from 189 patients and 105 NRAS mutations from 98 patients (Fig. 5A). Within these groups copy number alterations of the *LATS1* gene were observed with around 64% of sequenced BRAF mutant samples exhibiting heterozygous deletions and 2.8% of samples having homozygous deletions (Fig. 5B). Meanwhile almost 66% of sequenced NRAS mutant samples showed a shallow deletion of the *LATS1* gene, however no deep deletions were detected in these patients (Fig. 5B). Nevertheless, this early analysis strongly suggests that a majority of melanoma patients have alterations in *LATS1* expression.

**Figure 5.**
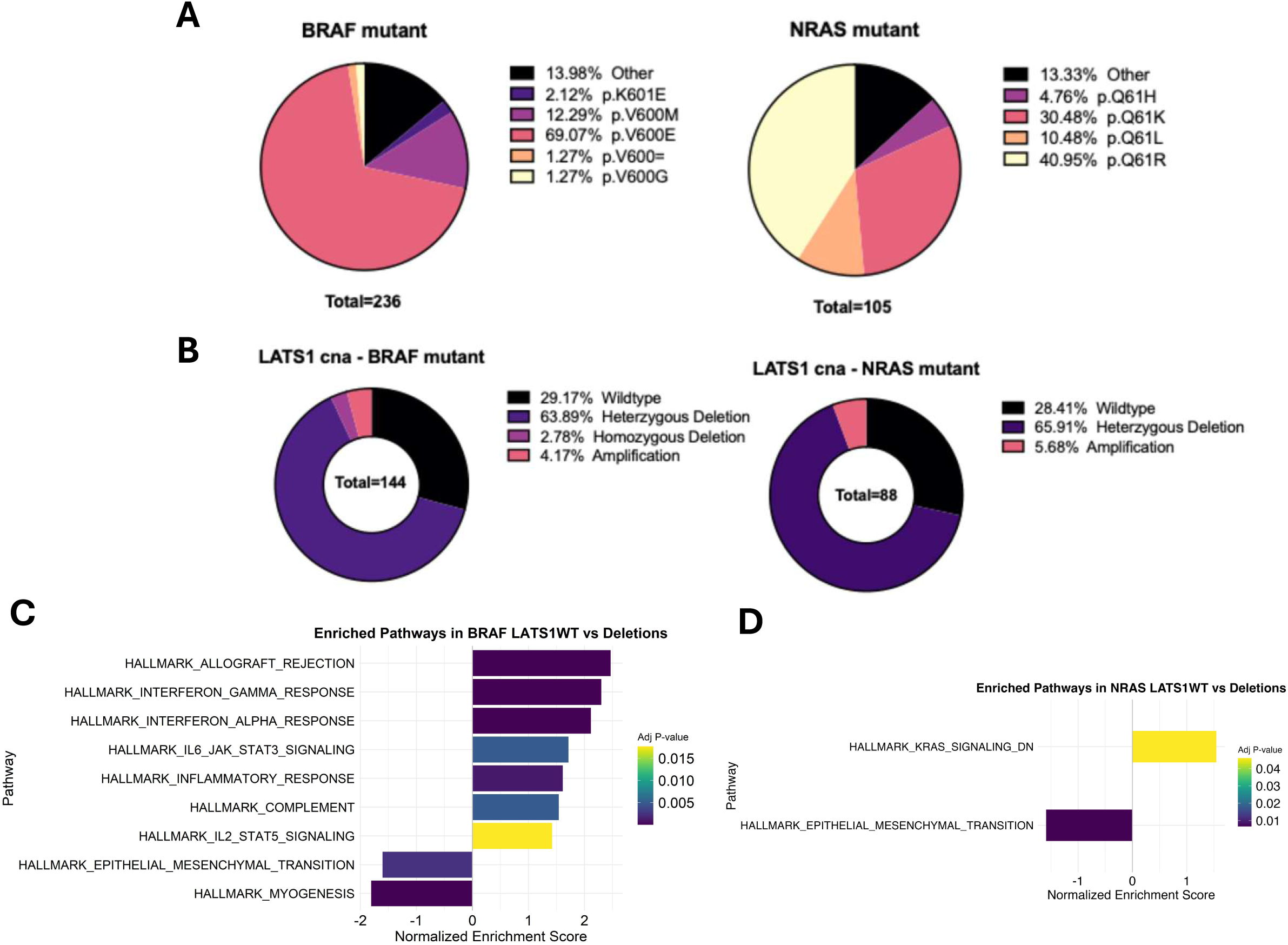
LATS1 expression correlates with immunotherapy response in melanoma patients and promotes interferon response in BRAF mutant melanoma. A) Pie charts representing the prevalence of BRAF and NRAS mutations in the Firehose legacy, TCGA cohort. B) Pie charts showing the percentage of patients with copy number alterations of the LATS1 gene within the BRAF mutant and NRAS mutant cohorts in the Firehose legacy dataset. C) Bi-directional bar plots showing GSEA normalized enrichment scores and the associated adjusted p-values for pathway enrichment analysis performed on RNA-seq data for patients with BRAF mutant melanoma and LATS1 wildtype (WT) expression or LATS1 deletions. D) Bi-directional bar plot of the normalized enrichment score (NES) and the adjusted p-value obtained from the GSEA Hallmarks database comparing patient with wildtype LATS1 expression and deletions in NRAS mutant melanoma.

To investigate the signalling changes associated with the differential *LATS1* expression, we utilised the RNAseq data in the Firehose Legacy cohort. First, we stratified patients by driver mutation (i.e. BRAF or NRAS), then further divided those groups into patients with no copy number alterations (cna) in the *LATS1* gene (WT) or those with heterozygous or homozygous *LATS1* deletions (Deletions). GSEA of these groups aligned with our previous zebrafish findings in the case of BRAF mutant melanoma. Here, we saw a positive enrichment in interferon-related and JAK/STAT pathways in the BRAF mutant *LATS1* WT group compared to the *LATS1* Deletions (Fig. 5C). However, comparing the NRAS mutant groups with respect to *LATS1* cna revealed very scarce differentially expressed genes with only 1 significantly positively and negatively enriched pathways resulting from the GSEA (Fig. 5D). We hypothesise that the lack of observed differences could be attributed to the high heterogeneity of NRAS mutant melanoma samples compared to the BRAF mutant samples as demonstrated by the high tumour mutational burden (Supplementary Fig. S5A). It is worth noting that the positively enriched pathway in the NRAS mutant *LATS1* WT group was the “down-regulation of KRAS signalling”, while EMT was negatively enriched in this group, suggesting less proliferative signalling in tumours presenting with *LATS1* WT expression (Fig. 5E).

To further explore the genes affected by changes in *LATS1* expression, we performed a correlation analysis between all genes and *LATS1* in both BRAF and NRAS mutant samples. From this analysis, we found that several matrix metalloprotease genes (MMPs) promoting melanoma metastasis were negatively correlated with *LATS1* expression in the NRAS mutant samples but not in the BRAF mutant tumours, suggesting higher invasiveness in the NRAS mutant cohort (Supplementary Fig. S5B and Fig. S5C). Moreover, we found that the expression levels of the DSBs marker H2AX negatively correlated with *LATS1* expression in both BRAF and NRAS mutant tumours (Supplementary Fig. S5B and Fig. S5C), albeit there was a higher correlation coefficient in NRAS mutant melanoma in line with the suppression of several DNA repair mechanisms observed in NRAS^Q61L^; *lats1^-/-^* zebrafish tumours. Finally, there was a positive correlation between levels of caspase 3 and 7 and *LATS1* expression in the BRAF mutant group (Supplementary Fig. S5C) and not the NRAS mutant group (Supplementary Fig. S5B). This is in line with our previous work showing that *LATS1* regulates apoptosis specifically in BRAF mutant cell lines (12). Taken together, these findings align with our zebrafish data, reinforcing the model’s relevance to human melanoma biology.

### *LATS1* promotes an anti-tumorigenic tumour microenvironment in human BRAF mutant melanoma

Given the previous observations, we raised the questions of whether *LATS1* i) regulates immune-related signalling pathways in BRAF mutant melanoma, ii) influences the tumour immune microenvironment (TiME), and finally, iii) whether its expression correlates with immunotherapy outcome in melanoma. Using the same *LATS1* WT vs. Deletions stratification in TCGA *BRAF*-mutant patients, we analysed IFN-γ signalling mediators. The data supported our zebrafish results and previous GSEA findings, showing significantly higher expression of *IFNGR1*, *IRF5*, and *IRF8* in *LATS1* WT tumours (Fig. 6A), along with increased expression of downstream JAK/STAT components *JAK1*, *JAK2*, and *STAT4* (Supplementary Fig. S6A) (24). We next examined MHC-I and MHC-II genes, which mediate antigen presentation and influence immune cell infiltration and immunotherapy response (29). Indeed, *B2M*, *HLA-A*, *CIITA*, and *HLA-DMA* were all upregulated in *LATS1* WT tumours (Fig. 6A; Supplementary Fig. S6A), alongside tumour-suppressive chemokines *CXCLS*, *CXCL10*, and *CXCL11* (30). These results suggest that *LATS1* wildtype expression induces an “immune-hot” TME in BRAF mutant human melanoma.

**Figure 6.**
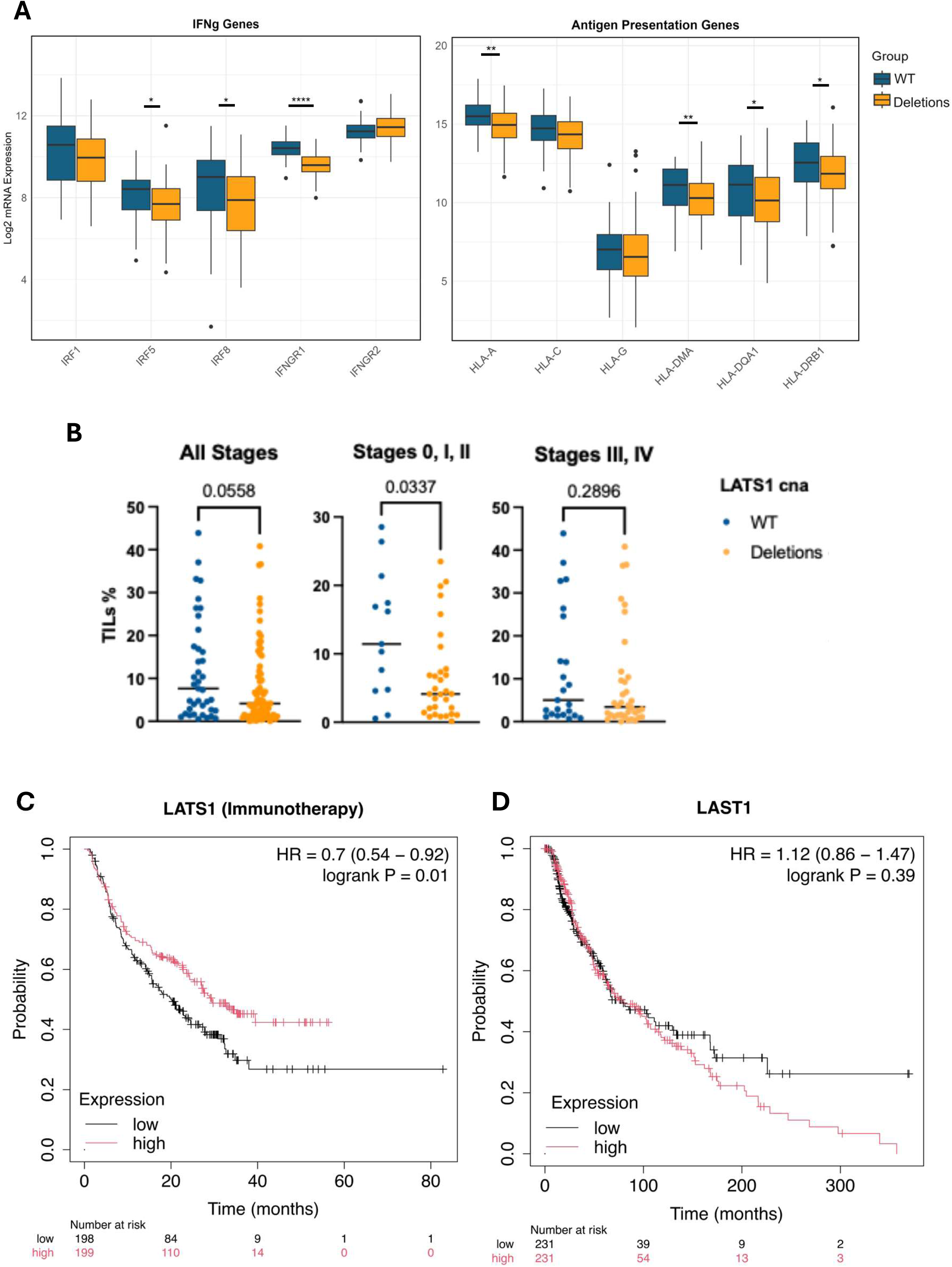
LATS1 wildtype expression promotes an anti-tumorigenic immune microenvironment. A) Boxplots for IFN-γ and antigen presentation related genes obtained the RNA-seq Firehose legacy dataset comparing patients with BRAF mutant melanoma with either *LATS1* wildtype expression (WT) or Deletions. Normal distribution was determined using the Shapiro-Wilk test, for normally distributed genes significance was calculated using an Unpaired t-test and for non-normally distributed genes significance was calculated using the Mann-Whitney test. *p< 0.05, **p<0.01, ****p<0.0001. B) Scatter plots showing the percentage of tumour-infiltrating lymphocytes in patients with *LATS1* wildtype expression (WT) or Deletions in BRAF mutant melanoma for the entire cohort of patients and when separated according to the disease staging. Statistical significance was calculated using the Mann-Whitney test. C) Kaplan-Meier curves and corresponding risk table comparing overall survival in melanoma patients with ’High’ and ‘Low’ *LATS1* expression treated with immunotherapy agents. The cut-off for group designation was the median *LATS1* expression. Significance was determined using the Log-rank test. D) Kaplan Maier curves and corresponding risk table showing the overall survival of patients included in the Firehose Legacy cohort with ‘High’ and ‘Low’ *LATS1* expression as determined by its median expression. Statistical analysis: Log-rank test.

This was further supported by applying the consensus TME deconvolution algorithm (31) to the patient RNAseq data to infer immune cell populations. Indeed, *LATS1* WT tumours showed higher infiltration of the tumour-suppressive CD4+, CD8+ T cells, and M1 macrophages, while *LATS1* deleted tumours had increased fibroblast infiltration, conventionally considered to be an immune-suppressive population (Supplementary Fig. S6B) (29). Moreover, no significant immune infiltration differences were inferred in NRAS mutant melanoma patients, which is in line with our observation of limited *LATS1-dependent* immune-related effects in zebrafish NRAS mutant melanoma (Supplementary Fig. S6C).

To validate this finding, we next used histological data generated by Saltz et al for the Firehose Legacy cohort (32). From our original cohort of patients with BRAF mutant melanoma, we analysed the percentage of tumour infiltrating lymphocyte (TILs) in 39 out of 42 patients in the *LATS1* WT group and 77 out of 96 patients with Deletions. We found a strong trend of higher TILs percentage in the *LATS1* WT group compared to the Deletions group (p=0.0558) (Fig. 6B). Further stratification by disease stage showed that those with early-stage melanoma (stages 0, I, II) had a significantly higher TILs percentage with *LATS1* WT expression compared to the Deletions group (p=0.0337) (Fig. 6B). However, in patients with late-stage melanoma (staged III, IV) there were no changes in TILs percentage between the groups (p=0.2896) (Fig. 6B). Conversely, when the same analysis was performed for patients with NRAS mutant melanoma, there were no significant changes in TILs percentage associated with the status of *LATS1* irrespective of the melanoma staging (Supplementary Fig. S6D).

TILs are positive predictors of immunotherapy response, therefore our next step was to test whether there are correlation of *LATS1* expression and survival in melanoma patients who received immunotherapy (29,33). To test this, we used KM plotter, which collects data from several databases, including those related to patients treated with immunotherapy agents (34). Indeed, the stratification of their melanoma cohort into high and low *LATS1* expression groups, showed a significantly improved overall survival in the *LATS1*-high group (Log-rank = 0.01) with immunotherapy treatment compared to the *LATS1*-low group (Fig. 5C). Worth noting that preferential survival is not associated with high *LATS1* expression in the patients of the TCGA cohort who were treated with other agents such as chemotherapy (Log-rank = 0.39) (Fig. 6D), confirming that the clinical significance of LATS1 seems to be immune-related.

Together, these findings identify a novel immune-regulatory function for *LATS1* in BRAF-mutant melanoma. Our results indicate that *LATS1* promotes IFN-γ signalling, antigen presentation, and recruitment of tumour-suppressive immune cells to the TME leading to a more favourable immunotherapy outcome, a mechanism first predicted from our zebrafish model and validated in human data.

## Discussion

This study demonstrates that *LATS1* influences melanoma progression through distinct, oncogene-dependent mechanisms, revealing novel regulatory roles with potential therapeutic implications. Using a zebrafish model, we show that *lats1* exerts a tumour-suppressive effect in BRAF^V600E^-driven melanoma, delaying tumour onset and improving tumour-free survival. In contrast, while *lats1* did not significantly alter tumour initiation in NRAS^Q61L^ melanoma, its loss was associated with a more invasive histological phenotype, including increased skin thickness and possibly brain invasion, suggesting a role in metastatic potential. We cannot determine from this study whether the brain tumours originated from brain melanocytes or if they metastasised from primary tumours; nevertheless, our results support a role for *lats1* in the appearance of melanoma in the brain. Interestingly, a recent study by Provance et al. linked YAP/TAZ activity to melanoma brain metastasis by mediating vascular mimicry and increasing the risk of brain metastasis (35). Therefore, this warrants further studies into the role of the MST2 pathway signalling in melanoma brain metastasis, which affects 50% of patients in the advanced stage of the disease and is associated with poor patient prognosis (20).

Molecular profiling of zebrafish tumours confirmed the model’s relevance for both BRAF and NRAS mutant melanoma. For BRAF mutant tumours, our data describes LATS1’s role in positively regulating immune response pathways, as evidenced by both pathway enrichment analysis and individual gene expression levels in the zebrafish model and the TCGA cohort. This finding is supported by data in other cancer types describing the emerging role of LATS1 in mediating immune response, including the regulation of the JAK/STAT signalling pathway downstream of interferon receptors (24,36). Further work from Zuo et al., showed that LATS1 was necessary for the antiviral innate immune response mediated by the activation of the type-I interferon response by promoting STAT1 phosphorylation (13). Similarly, another study described the role of LATS1 in activating STAT1 to promote interferon-gamma activation and immune cell infiltration in endometrial cancer (14). Conversely, Moroishi et al. demonstrated that the dual loss of LATS1 and LATS2 expression correlated with the induction of an anti-tumorigenic immune response in melanoma mouse xenograft models, indicating that these two tumour suppressors might have different functions in the modulation of immune responses (37). To the best of our knowledge, this is the first study that directly correlates LATS1 expression with a favourable immunotherapy response in melanoma. Therefore, the zebrafish tumour data and the analysis of human melanoma cohorts support the role of LATS1 as an orchestrator of an anti-tumorigenic immune response in BRAF mutant melanoma and provide a mechanistic basis for this effect. It is worth noting that our animals are complete *lats1* knockouts, so we cannot rule out the possibility that part of this effect could be due to the lack of expression of *lats1* in the immune system. However, the fact that NRAS^Q61L^*; lats1^-/-^* tumours do not show the same phenotype strengthens our hypothesis. Moreover, in human tumours, the immune cells are not defective for LATS1. Therefore, we conclude that the role of the tumour suppressive effect of LATS1 in BRAF-driven tumours is, most likely, due to LATS1 signalling in tumour cells.

A distinct molecular profile was seen in the NRAS^Q61L^ zebrafish tumours with loss of *lats1*, exhibiting a more invasive signature and showing upregulation of DNA damage repair (DDR) transcriptional programmes, which could allow the tumour to overcome genomic instability and, potentially limit the cytotoxic response to chemotherapeutics (38). Specifically, we saw higher Fanconi Anaemia pathway enrichment in NRAS^Q61L^*;lats1^-/-^*. While in other cancer types FA pathway is downregulated, in melanoma FA is upregulated and more so in metastatic lesions (26,38). Interestingly, high FA genes have been previously correlated with increased Breslow depth (26). This is evident in our histological analysis showing increased thickness in NRAS^Q61L^*; lats1^-/-^* samples, which could be mediated in part by the elevated DDR in this group. Furthermore, increased migration gene markers were seen in the NRAS^Q61L^ zebrafish tumours with loss of *lats1*, which correlates nicely with the elevation of MMPs in patient samples in the same conditions. MMPs are well-described to degrade the extracellular matrix, promoting tumour migration and metastasis (39). The regulation of MMPs by LATS1 has been previously described in other cancer types (40–42). Therefore, findings from our study warrant further investigations into this relationship in the context of NRAS mutant melanoma and the possible therapeutic benefit of using MMP inhibitors in this specific patient cohort. Meanwhile, the regulation of the cell cycle and DNA damage pathways are well-described roles for LATS1 (43). This includes both YAP-independent and YAP-dependent mechanisms that result in G2/M transition blockade and the induction of apoptosis in response to DNA damage (43). Our data suggest that patients with NRAS mutant melanoma and LATS1 wildtype expression might benefit from treatments such as PARP inhibition, as the expression profile of these tumours suggests an impaired DDR capacity, which could increase their susceptibility to this treatment..

In summary, our zebrafish model reveals distinct tumour-suppressive roles for LATS1 in BRAF and NRAS-driven melanoma. These findings have translational relevance, suggesting LATS1 status could inform immunotherapy stratification in BRAF-mutant patients and highlight potential synthetic lethality strategies in NRAS-driven melanoma.

## Materials and Methods

### Ethical approval and Project authorisation

All experiments carried out were granted ethical approvals by the Animal Research Ethics Committee in UCD, under the following reference numbers: 18-18-Halasz and 18-26-Gomez. The project authorisation was granted by the Health Products Regulatory Authority of Ireland under the reference number AE18982/P148.

### Zebrafish husbandry

Zebrafish were maintained at a 14-hour light and 10-hour dark cycle in a recirculating water system set at 28°C. Colonies were maintained at a density of 3-5 fish/litre. Zebrafish colonies were maintained under 2 years of age. The fish were fed with the recommended zebrafish diet consisting of brine shrimps and ZM300 pellets.

### Generation of a zebrafish melanoma model

The melanoma model was generated *in mitfa* mutant (nacre)/*tp53* mutant (M214K) zebrafish. CRISPR/Cas9 technology was used to generate the lats1 knockout strain. Heterozygous *lats1^+/-^* zebrafish generated from a founder generation were cross-bred to achieve a homozygous *lats1^-/-^* strain. gRNAs used were designed using https://chopchop.cbu.uib.no and the sequence details are included in the supplementary materials. The introduction of the oncogene was done using Tol-2 mediated transgenesis of a miniCoopR plasmid harbouring either BRAF^V600E^ or NRAS^Q61L^ or a GFP control. The plasmids were generated as previously described (17). Zebrafish were spawned and the embryos were collected within 10 minutes to allow for embryo fertilisation. 25 pg of miniCoopR plasmid and 25 pg of Tol2 transposase mRNA were injected into the single-cell stage embryo using a microinjector (Nanoject III, Drummond). The injected embryos were incubated at 28°C, and only larvae with similar melanocyte rescue efficiency were moved to tanks at 131 hours post-fertilisation. Transgenic fish were monitored twice daily up to 12 days post-fertilisation and then monitored for welfare scoring and tumour development daily. Zebrafish showing any signs of compromised welfare or tumours that occupied more than 20% of the body surface were immediately culled using 0.6 mg/ml Tricaine (Sigma-Aldrich, Burlington MO, USA).

### Zebrafish tumour development and survival analysis

Zebrafish were monitored daily, and the number of tumours and tumour location per fish were recorded. Tumour-free survival analysis was performed in R Studio (v 2023.06.02) using the Survival package (v 3.5-7). The Survminer package (v 0.4.9) was used to plot the Kaplan-Meier curves. The statistical significance of the survival curves was calculated using the Log-rank test. Tumour numbers recorded were analysed in R Studio (v 2023.06.02) and were correlated with the zebrafish genetic background using a logistic regression model using the Glmnet package (v 4.1-8). The zebrafish sample numbers per group were determined by performing power calculations to ensure statistical significance. The sample size was determined to be 20 animals per group.

### Histological analysis of zebrafish sections

Zebrafish were euthanised using Tricaine (Sigma-Aldrich,), placed on ice and the whole body was fixed in 10% neutral buffer formalin (4% Formaldehyde (Sigma-Aldrich), 0.4%, Sodium phosphate monobasic (Sigma-Aldrich), 0.65%, Sodium phosphate dibasic Sodium phosphate dibasic, 1.5% Methanol (Sigma-Aldrich), 93% water) for 24h, followed by incubation in 0.35 M EDTA (Promega, Madison WI, USA) for 7 days for bone decalcification. The zebrafish were either cut transversely or longitudinally. Tissue samples were subsequently processed, paraffin-embedded and 5 µm sections were stained with haematoxylin and eosin (Sigma-Aldrich) (HCE). Epidermal and scale measurements overlying the dorsal spinous process, just rostral to the dorsal fin, were performed by Dr Pamela Kelly, a board-certified veterinary pathologist (European College of Veterinary Pathologists), using an Olympus SC50 camera-equipped microscope and CellSens image acquisition software.

### Protein and RNA extraction from zebrafish tumours

Protein and RNA were extracted from matching samples by dissecting the tumour in half. One half was used for protein extraction using RIPA lysis buffer (50 mM Tris-HCl (Sigma-Aldrich) pH 7.4, 150 mM NaCl (Fisher Scientific, Waltham MA USA), 1 mM EDTA, 0.1% SDS (Fisher Scientific), 1% NP-40 (Calbiochem, Rahway NJ, USA), 0.5% Sodium Deoxycholate (Sigma-Aldrich) supplemented with protease and phosphatase inhibitors (Roche, Basel, Switzerland). The tumour tissue was disrupted using a homogeniser. The lysate was centrifuged at 12000 RCF for 10 minutes at 4°C, and the supernatant was transferred to a new labelled tube. The pellet was further lysed using 8M Urea (Thermo Scientific)/50 mM Tris-HCL pH 7.5, supplemented with protease and phosphatase inhibitors. The samples were sonicated (Ultrasonic Homogeniser SKL-150W, Syclon) at 20% power 3x for 10 seconds and centrifuged at 18000 g for 10 minutes at 4°C. The supernatant was collected and added to that from the initial lysis step. RNA was extracted from the second half of the tumour using the RNeasy Plus Mini Kit (Qiagen, Hilden, Germany). Briefly, RLT lysis buffer was added to the tumour and the tissue was disrupted using a homogeniser, followed by sonication (Ultrasonic Homogeniser SKL-150W, Syclon) at 20% power 3x for 10 seconds. The remaining RNA extraction steps were performed following the manufacturer’s protocol.

### SDS-PAGE and Western blots

Protein lysates were resolved using denaturing sodium dodecyl sulphate polyacrylamide gels (SDS-PAGE) in running buffer (25 mM TrisBase (Merck, Rahway NJ, USA), 190 mM Glycine (Sigma-Aldrich), 0.1% SDS. The gels were equilibrated in transfer buffer (25 mM Tris, 190 mM Glycine, 20% Methanol) for 10 minutes. The proteins were then transferred onto polyvinylidene fluoride (PVDF) membranes (Millipore, Burlington, MA, USA) using the BioRad Transblot electrophoretic semi-dry transfer cell system. Membranes were then blocked with 4% Bovine Serum Albumin (BSA, Sigma-Aldrich) in TBST (50 mM Tris, 150 mM NaCl, 0.1% Tween 20 (Sigma-Aldrich) for 1 hour at room temperature and incubated with the primary antibody overnight at 4°C. The following primary antibodies were used, from Cell Signalling Technology (Danvers,, MA, USA) : LATS1 (C66B5) (3477), YAP (D8H1X) XP (14074), β-Actin (13E5) (4970); from Santa Cruz (Dallas, TX, USA): N-Ras (C-20) (sc-519), α-Tubulin (sc-8035); from Sigma-Aldrich: ERK1/2 (M5670), Phospho-ERK1/2 (M8159); from LS Bio (Newark, CA, USA): B-Raf (LS-C353922). The following horseradish peroxidase (HRP)-conjugated secondary antibodies from Cell Signalling Technology were used to visualise the proteins by chemiluminescence: Anti-Rabbit IgG (7074), Anti-Mouse IgG (7076). The uncropped western blot images are available in the supplementary materials file.

### RNA-sequencing for zebrafish tumours and differential expression analysis

RNA-sequencing was performed by Biomarker Technologies (BMK) GMBH (Munster, Germany) using the Illumina Novaseq 6000 system. The quality of the sequencing data received was assessed using FastQC prior to any downstream analysis. Processing of the mRNA reads was performed using the RNAlysis software (44) and pseudoalignment and quantification of the reads were done using the Kallisto package against the zebrafish genome (Danio_rerio.GRCz11.110). The raw expression matrix after filtration was processed using the DESeq2 package in R Studio (v 2023.06.02) to generate the differentially regulated genes and to generate the normalised expression matrix, which was then Log_2_ transformed for individual gene expression analysis.

### Proteomics sample preparation

Sample clean-up was performed using the SP3 protocol (45). Protein extracts were quantified with Pierce™ Bradford Plus Protein Assay Reagent (Thermo Scientific), and the samples were normalised to 300 µg of protein. DNA and RNA were removed from the samples using Benzonase Nuclease (Sigma-Aldrich) degradation. Samples were reduced and alkylated using 10 mM dithiothreitol (DTT, Sigma-Aldrich) and 20 mM iodoacetamide (IAA, Sigma-Aldrich), respectively. Acetonitrile (Sigma-Aldrich) was added to a final concentration of 70%, followed by the addition of hydrophobic and hydrophilic Speedbead magnetic Carboxylate Modified Particles (GE Healthcare, Chicago IL, USA). The samples were rotated at room temperature for 1 hour. Samples were placed on a Dynamag™ stand, and the supernatant was aspirated. The beads were washed with 70% ethanol and 100% acetonitrile. Finally, the beads were resuspended in 50 µl of 50 mM ammonium bicarbonate (Sigma-Aldrich) before trypsin digestion. Protein digestion was performed using Sequencing Grade Modified Trypsin (Promega) at a concentration of 1 µl trypsin to 50 µg protein, and samples were incubated overnight on a thermomixer (Fisher) at 500 RPM at 37°C. The following day, 10 µl of beads and 1 ml of 100% acetonitrile were added to each sample, and the samples were incubated on a rotator for 1 hour. The supernatant was discarded, and the beads were washed with 100% acetonitrile. Peptides were eluted from the beads with MS-grade H_2_O (Fisher Scientific), and 2 µl of 10% acetic acid (Sigma-Aldrich) was added to each sample. The samples were vacuum centrifuged using CentriVap Concentrator (Labconco) and resuspended in 0.1% trifluoroacetic acid (Sigma-Aldrich) for mass spectrometry analysis.

### Mass spectrometry data acquisition

Digested peptides were run on a Bruker timsTof Pro mass spectrometer (Bruker) connected to an Evosep One liquid chromatography system (Evosep One) using Evosep tips. The autosampler was configured to pick up each tip, elute and separate the peptides using a set chromatography method (46). The mass spectrometer was operated in positive ion mode with a capillary voltage of 1750 V, dry gas flow of 3 l/min and a dry temperature of 250°C. All data were acquired with the instrument operating in trapped ion mobility spectrometry (TIMS) mode. Trapped ions were selected for ms/ms using parallel accumulation serial fragmentation (PASEF). A scan range of 100-1700 m/z was performed at a rate of 5 PASEF MS/MS frames to 1 MS scan with a cycle time of 1.03 seconds (47). Chromatography Buffers used: Buffer A: 99.9% water, 0.1% formic acid, Buffer B: 99.9% acetonitrile, 0.1% formic acid (Sigma-Aldrich, 33015-M. All solvents are LC/MS grade.

### Fragpipe software

The raw proteomic data were searched against the *Danio rerio* subset of the UniProt SwissProt database (unreviewed) using the Fragpipe proteomics pipeline (v 20.0) with specific parameters for trapped ion mobility spectra data-dependent acquisition (TIMS DDA). The LFQ-MBR workflow was selected. Details include MS Fraggers parameters: precursor mass tolerance +/-20ppm, fragment mass tolerance +/-20ppm, strict trypsin, fixed modifications oxidation of methionine and N-terminal acetylation.

### Analysis of proteomics data

Label-free quantification values from proteomics data were acquired, and missing data were imputed using a mixed imputation approach, where the missing values were replaced by the mean value of the group when the missing values were present in less than 80% of the samples. The data was then log2-transformed, and individual protein expression was plotted using Graphpad Prism Software (v 10).

### Pathway enrichment analysis for transcriptomics data

Gene set enrichment analysis (GSEA) was performed on the ranked list of all genes using the fgsea package. The Biomart package (v 2.56.1) was used to convert gene names to Ensembl IDs using the “drerio_gene_ensembl” database. Pathways with a p-adjusted value <0.05 were considered to be significant.

### Dataset integration and analysis

The bulk RNA-seq and proteomics datasets were integrated using the Multiset correlation and factor analysis (MCFA) method with default parameters (23). Genes identified in factors that explain the most variance across the experimental conditions were converted to the human orthologs using the orthogene package in R studio (v 1.6.1). Enrichment analysis was performed using Gene Set Enrichment Analysis (GSEA) to identify significantly upregulated or downregulated pathways in our dataset. Pathways with a p-adjusted value <0.05 were considered to be significant.

### TCGA data analysis

Transcriptomics, gene mutational data and clinical data for the SKCM TCGA Firehose legacy dataset were downloaded from cBioPortal. Kaplan-Meier curves were generated using the KM plotter online tool (34). Differential expression analysis was performed using the DESeq2 package (v 1.40.2). Gene set enrichment analysis was performed using the fgsea package (v 1.26.0). The consensus tumour microenvironment (consensusTME) cell estimation method was used to deconvolve the TME of TCGA samples (31), employing gene sets specific to human skin cutaneous melanoma (SKCM). Differentially abundant cell types were identified by min-max scaling the inferred cell abundances. Median normalised enrichment scores (NES) were computed for each patient group, and differences between groups were quantified as the log₂ ratio of median NES values. Statistical significance was assessed using the Wilcoxon rank-sum test with Benjamini–Hochberg correction. An FDR threshold of 0.05 was applied.

### Statistical analysis

Statistical analyses were carried out using GraphPad Prism Software (v 10) or in R Studio (v 2023.06.02). All statistical tests were done on a minimum of 3 biological repeats. Normal distribution of the data was first determined using the Shapiro-Wilk test prior to further statistical analysis. The specific tests used are indicated in the associated figure legends. A cut-off value of p <0.05 was considered to be statistically significant for all analyses performed.

**Table 1.** H&E section for tumour-bearing zebrafish. The genetic background and the number of transverse sections analysed per group are indicated in the first column. Zebrafish sections that show evidence of melanoma invasion into the muscles are highlighted in red box while sections showing melanoma invasion into the brain are highlighted in a blue box.

| Transgenic line<br>(Total # of fish) | # with muscle invasion | # with invasion to spinal cord | # with brain invasion | H&E images |
|---|---|---|---|---|
| ***Tg: mitfa:BRAF<sup>V600E</sup>; P53<sup>mut</sup>;lats1<sup>+/+</sup>***<br>(3) | 1 | 1 | 0 |  |
| ***Tg: mitfa:BRAF<sup>V600E</sup>; P53<sup>mut</sup>;lats1<sup>-/-</sup>***<br>(4) | 4 | 3 | 0 |  |
| ***Tg: mitfa:NRAS<sup>Q61L</sup>; P53<sup>mut</sup>;lats1<sup>+/+</sup>***<br>(4) | 4 | 4 | 0 |  |
| ***Tg: mitfa:NRAS<sup>Q61L</sup>; P53<sup>mut</sup>;lats1<sup>-/-</sup>***<br>(5) | 5 | 5 | 3 |  |

## Acknowledgements

We want to thank Amaya García Muñoz and Philip Cotter for technical support. We thank the support from UCD Biomedical facilities/Conway zebrafish facility, and in particular, Mark A. Dalton and Stephen Reynolds. We thank Darrel Andrews, Alison Reynolds, Tess McCann, Brendan Kennedy and Yolanda Álvarez for advice and support in the work with zebrafish. We are grateful to all the patients who donated the samples used in this study.

## Data availability statement

The mass spectrometry proteomics data have been deposited in the ProteomeXchange Consortium via the PRIDE (48) partner repository. The raw and processed RNA-seq files have been deposited in NCBI’s Gene Expression Omnibus (49). The deposited datasets will be released and accession numbers disclosed upon publication of the article. Prior to publication, the data are available from the corresponding authors upon reasonable request.

## Author contribution

Conceptualization: D.M.; Data curation, N.A., L.G.G. and D.M.; Formal analysis, N.A., L.G.G., P.K., SP, BC, E.M.T, and D.M.; Funding acquisition, W.K., N.A and D.M.; Investigation, N.A. L.G.G., S.S., E.O. and D.M.; Methodology, N.A., L.I.M., D.E., P.K., E.M.T., A.B, A.H., S.P., B.C., K.W. and D.M.; Project administration, M.H., P.K., A.H., N.A., L.G.G. and D.M.; Resources, D.M.; Supervision, M.H., A.H. and D.M.; Visualisation, N.A., D.E., L.I.M., and D.M.; Writing—original draft, N.A. L.G.G. and D.M.; Writing—review and editing, N.A., P.K. A.H., LIM, D.E., W.K., K.W. and D.M. All authors have read and agreed to the published version of the manuscript.

## Ethical approval by the Institutional Review Board Statement

All experiments carried out were granted ethical approval by the Animal Research Ethics Committee in UCD, under the following reference numbers: 18-18-Halasz and 18-26-Gomez.

---

**Supplementary figure S1.**
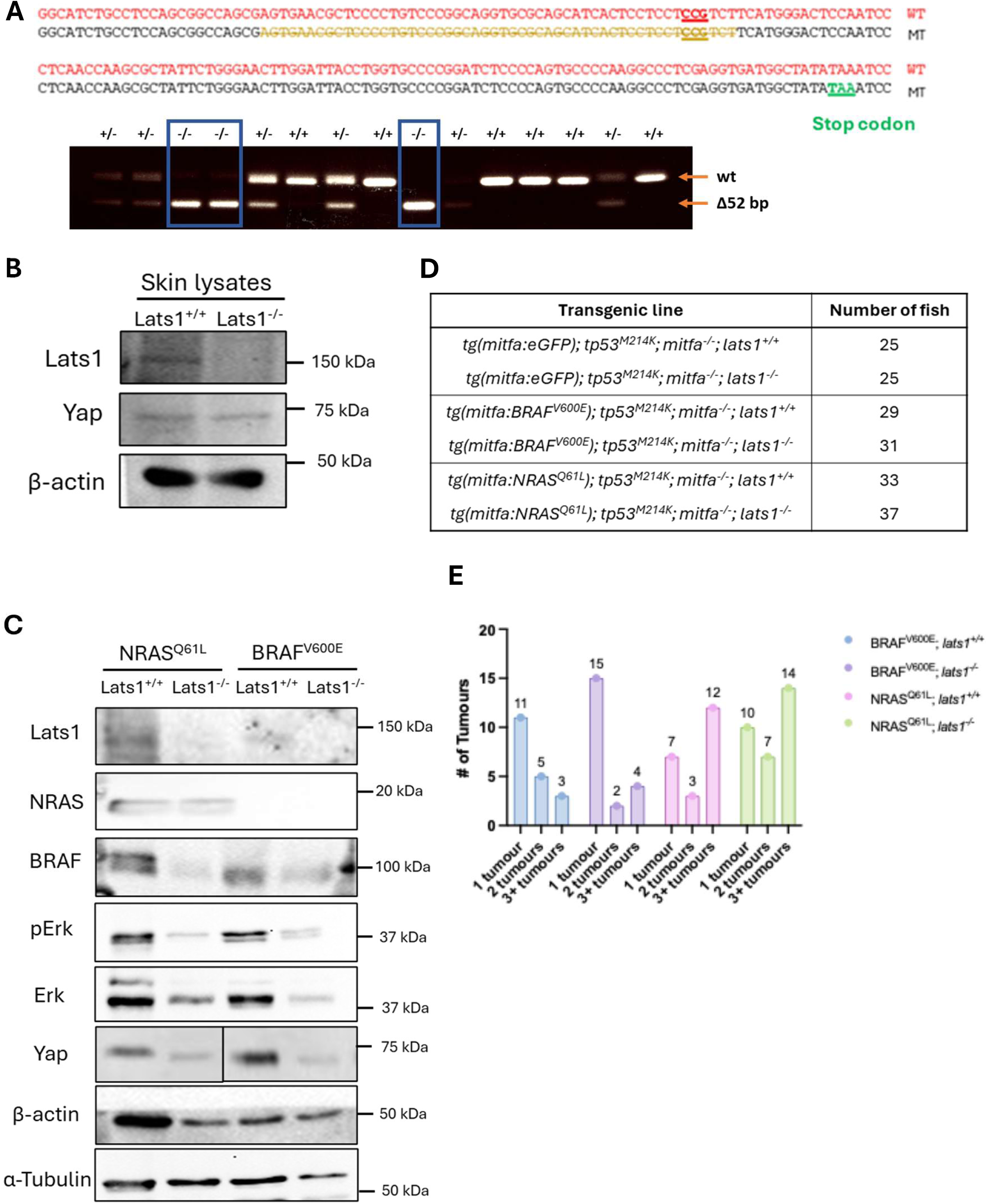
A) Sequence for *lats1* gene with the generated stop codon and associated PCR agarose gel confirming the generation of a knockout strain. B) Western blots of zebrafish skin lysates from the control groups. C) Western blots of zebrafish fish isolated tumour lysates. D) Table showing the number of zebrafish generated per genetic group. E) Bar-plot showing the number of fish per genetic group that developed either 1 tumour, 2 tumours or, 3+ tumours.

**Supplementary figure S2.**
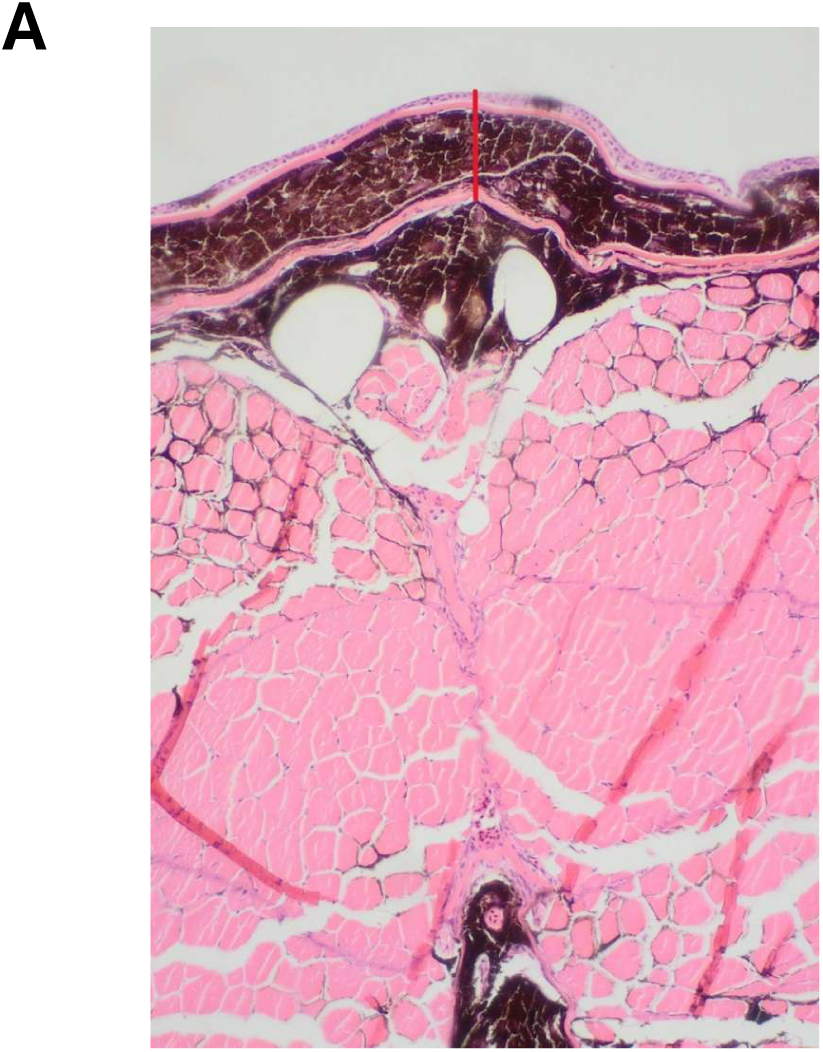
A) Histological image showing the epidermis, scale, dermis and muscle overlying the dorsal spinous process (Hematoxylin and eosin, 10X). The red line represents the skin measurement used for histological statistical analysis.

**Supplementary figure S3.**
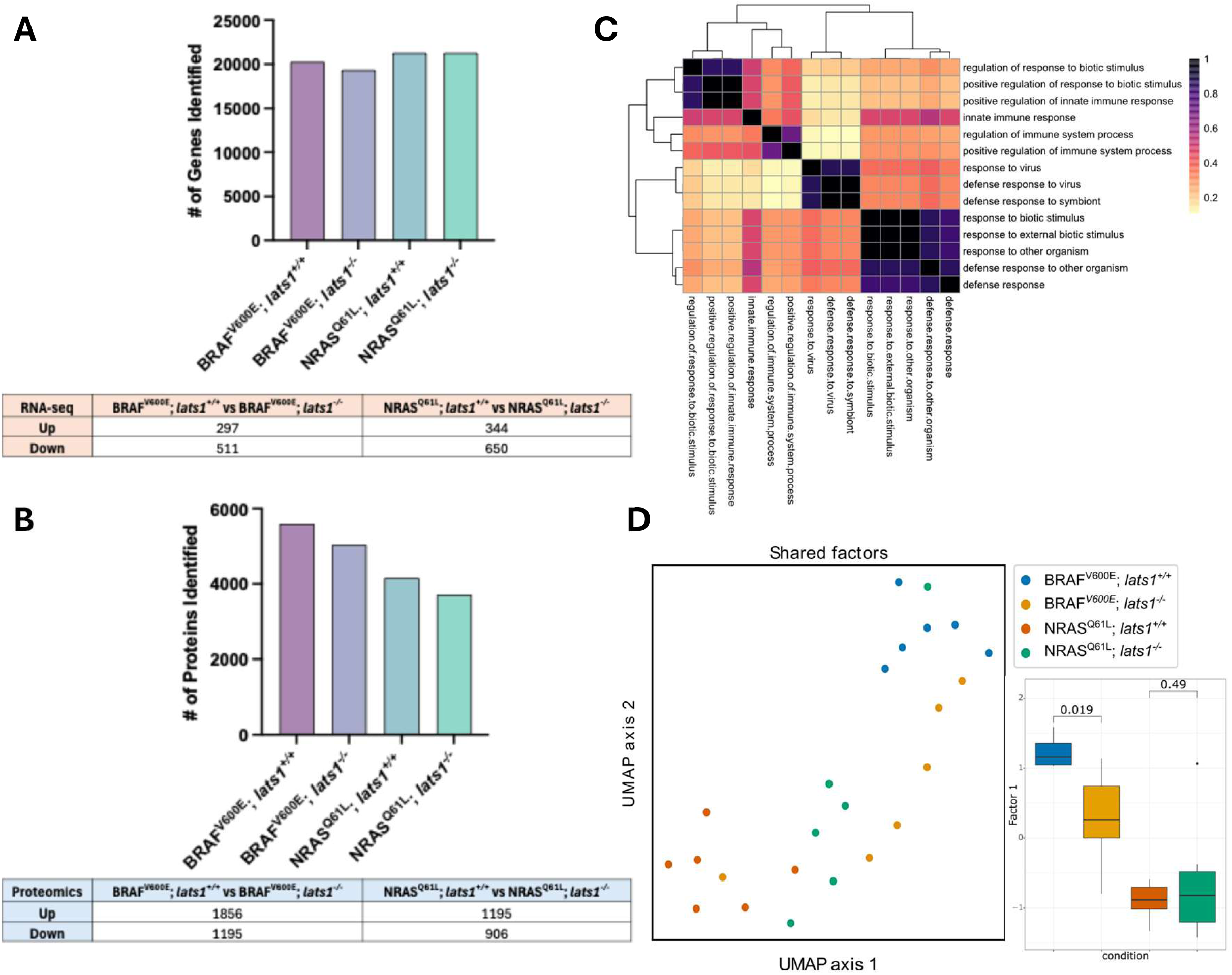
(A, B) Bar-plots and associated tables shows the number of genes (A) and proteins (B) identified in the raw data files for each genetic group. C)Heatmap illustrating pairwise pathway similarities calculated using the Jaccard similarity index for immune-related pathways identified using GSEA for GO BP of RNA-seq data of BRAF^V600E^ samples. Darker shades represent higher similarity values. D) UMAP showing the shared factors generated by multi-omics data integration and the clustering of the different samples with the associated box-plots showing the significance of factor 1 which is contributing towards the variances of the BRAF^V600E^ melanoma groups.

**Supplementary figure S4.**
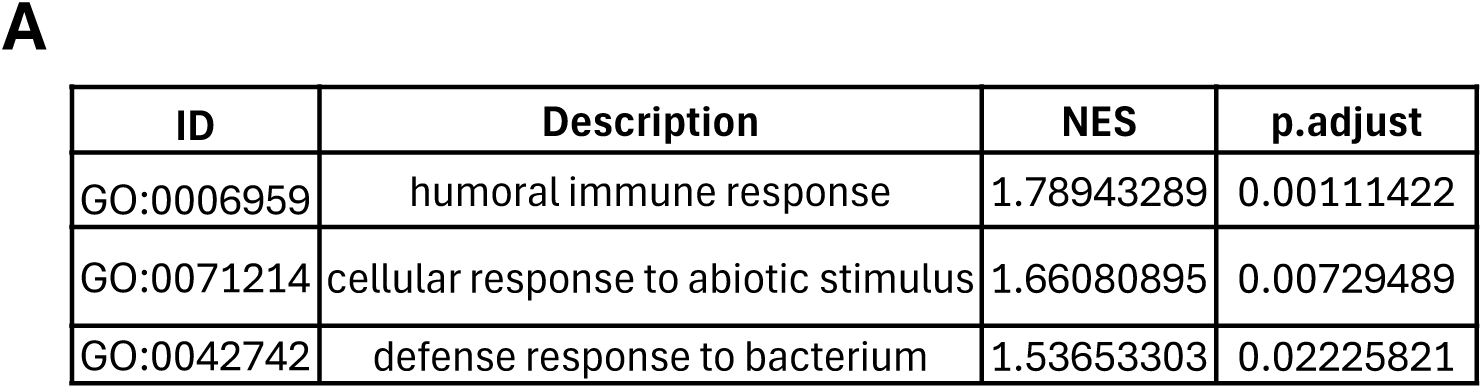
A)Table of the immune-related pathways identified in GSEA GO BP analysis of the NRAS^Q61L^ tumours along with the normalized enrichment scores (NES) and the p adjusted values.

**Supplementary figure S5.**
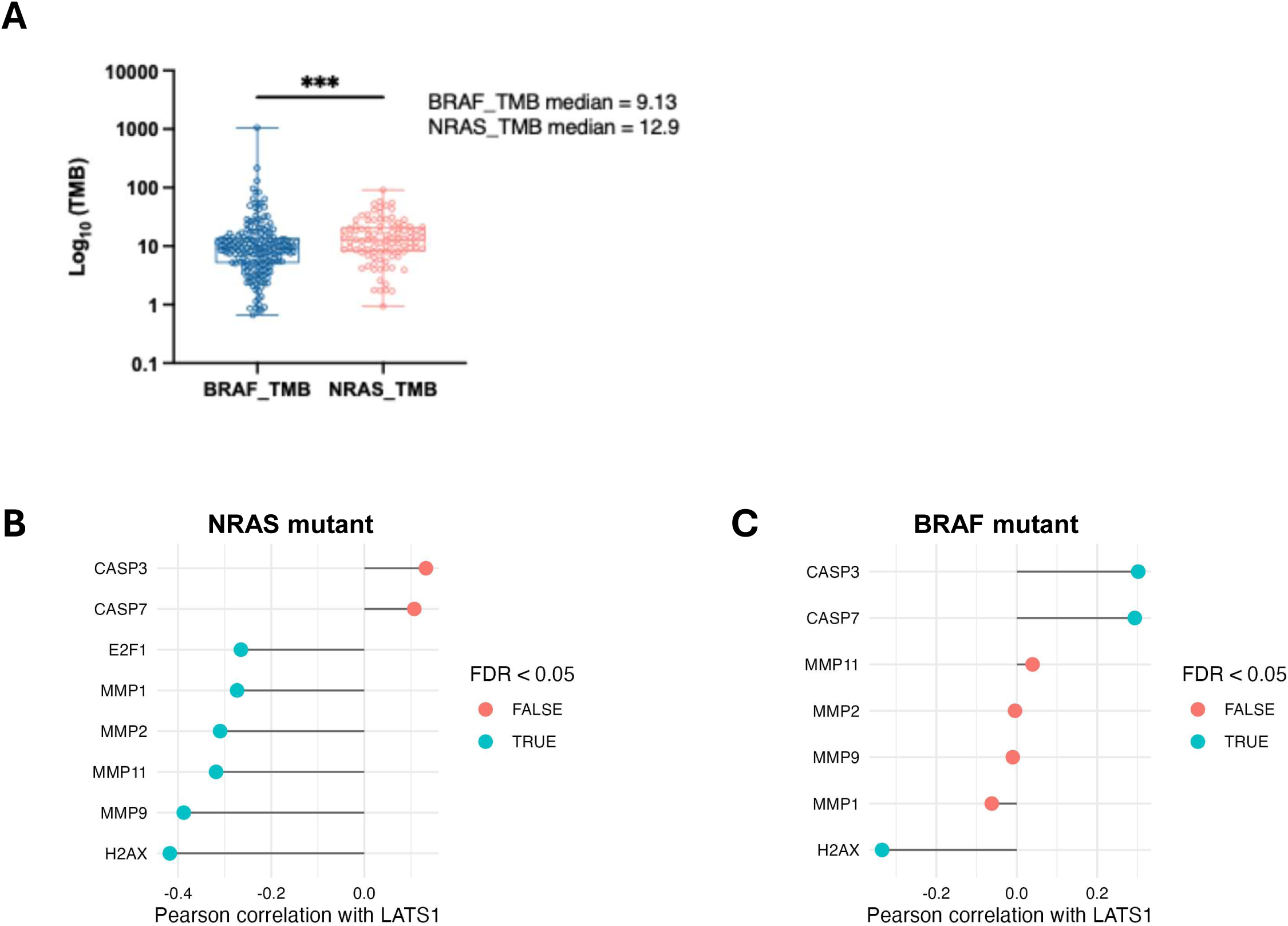
A) Box-plots showing comparing the tumour mutational burden (TMB) of patients with BRAF and NRAS mutant melanoma. Statistical significance was calculated using the Mann-Whitney test. ***p<0.001. (B, C) Lollipop plots of genes correlating with the *LATS1* expression with either NRAS (B) or BRAF (C) mutant melanoma. Correlation values were determined using Pearson correlation and genes with FDR <0.05 were considered to be significantly correlated and marked in red. Genes with FDR >0.05 were not significantly correlated and were marked in red.

**Supplementary figure S6.**
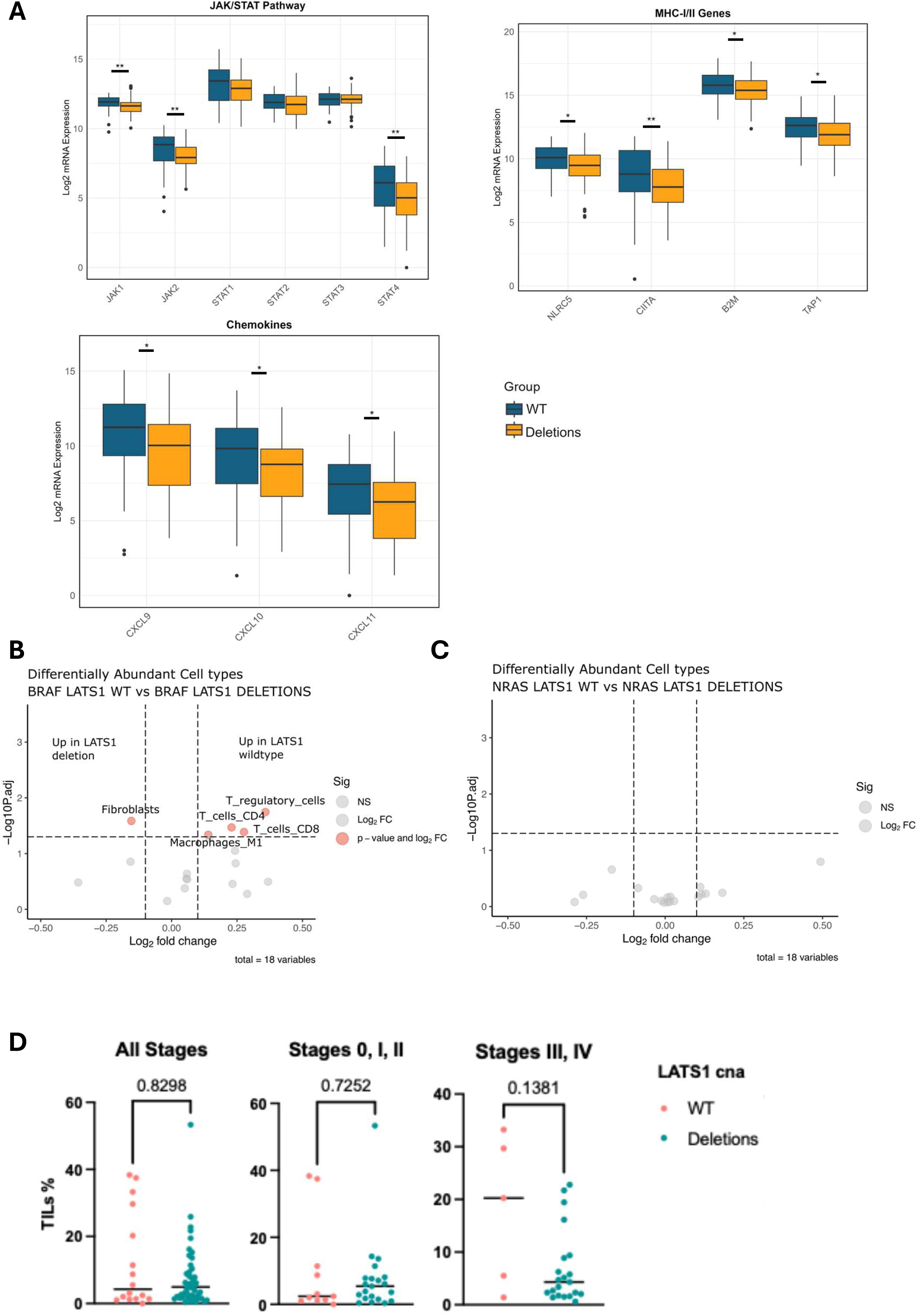
A) Box-plots for antigen presentation, JAK/STAT and pro-inflammatory chemokines related genes obtained the RNA-seq Firehose Legacy dataset for patients with BRAF mutant melanoma with either *LATS1* wildtype expression (WT) or Deletions. Normal distribution was assessed using the Shapiro-Wilk test, for normally distributed genes significance was calculated using an Unpaired t-test and for non-normally distributed genes significance was calculated using the Mann-Whitney test. *p< 0.05, **p<0.01. B) Volcano plot showing significantly differentially abundant immune cells detected using the TME consensus RNA-seq deconvoluting algorithm in BRAF mutant melanoma when comparing *LATS1* wildtype expression (WT) with Deletions. C) Volcano plot showing differential abundance of immune cells detected using the TME consensus RNA-seq deconvoluting algorithm for NRAS mutant melanoma when comparing *LATS1* wildtype expression (WT) with Deletions. D) Scatter plots showing the percentage of tumour-infiltrating lymphocytes in patients with *LATS1* wildtype expression (WT) or Deletions in NRAS mutant melanoma for the entire cohort of patients, and when separated according to the disease staging. Statistical significance was calculated using the Mann-Whitney test.

## Supplementary materials

### Danio rerio lats1 gene sequence and gRNA primer design for lats1 knockout

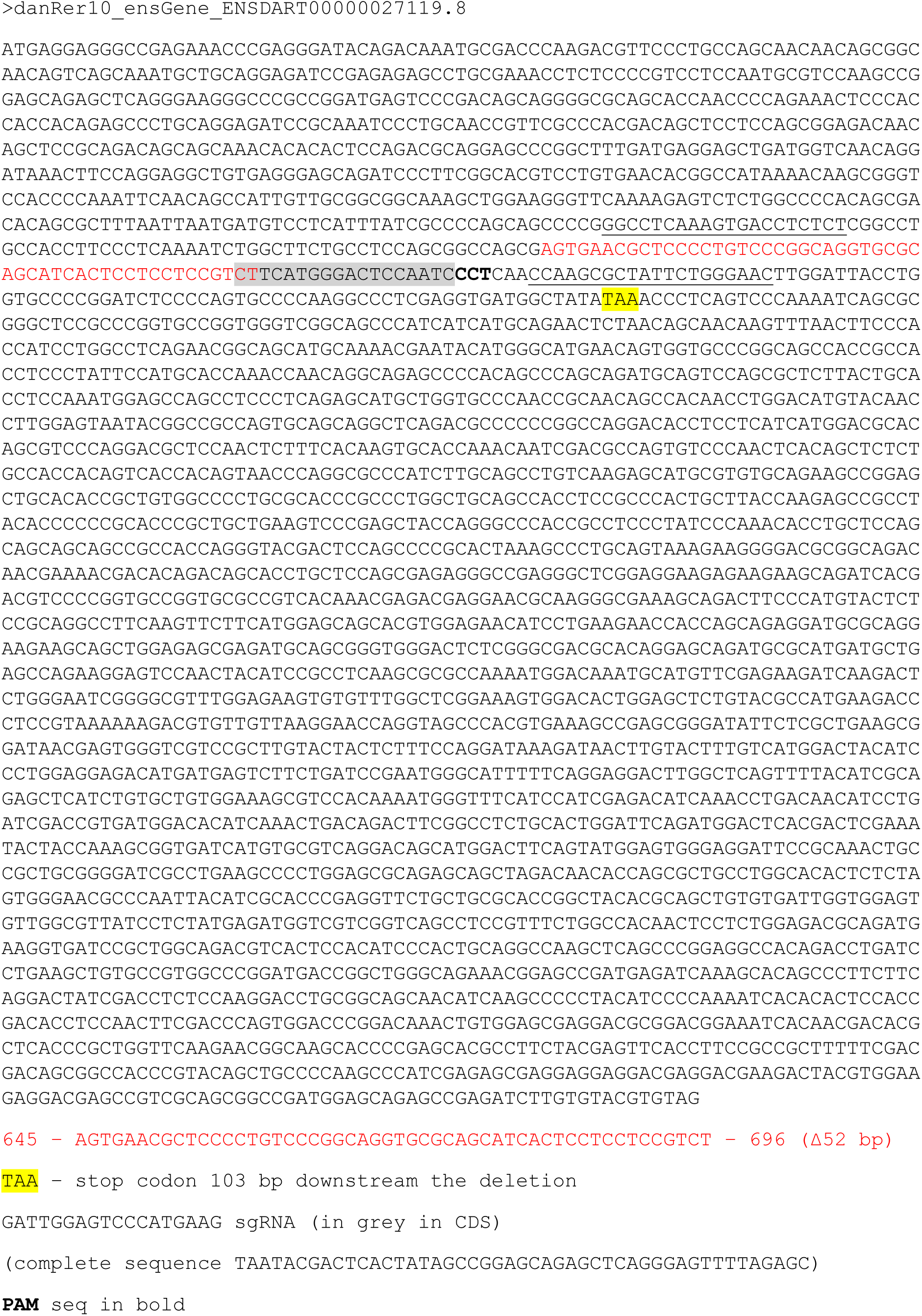
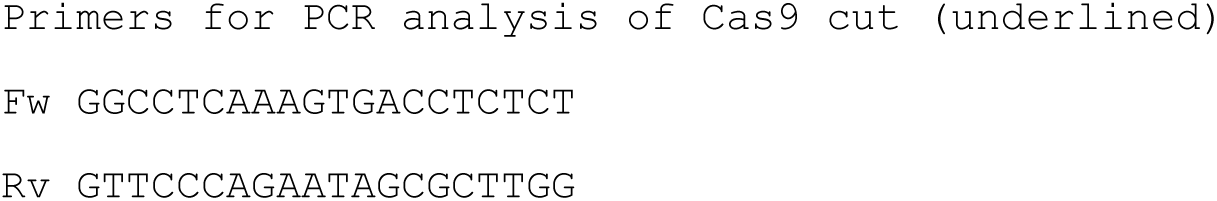

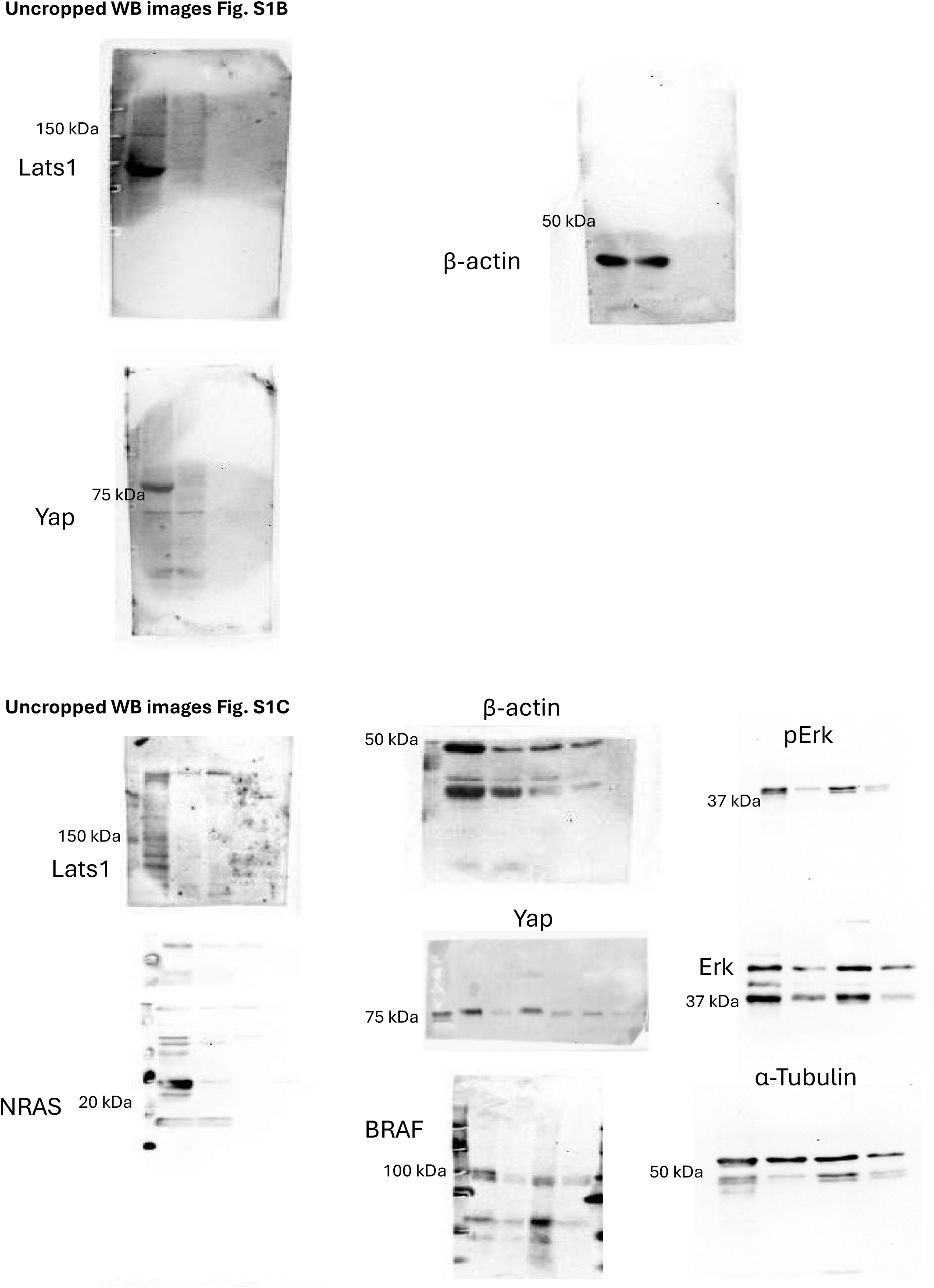

